# Differential side-effects of *Bacillus thuringiensis* bioinsecticide on non-target *Drosophila* flies

**DOI:** 10.1101/541847

**Authors:** Aurélie Babin, Marie-Paule Nawrot-Esposito, Armel Gallet, Jean-Luc Gatti, Marylène Poirié

## Abstract

Biopesticides based on *Bacillus thuringiensis* (*Bt*) spores and toxins are alternate pest management solutions widely used to control insect pests. Their increasing use could lead to accumulation in the environment, hence leading to chronic exposure of non-target organisms. Here, we tested for potential non-intentional side-effects of chronic exposure to *Bt* biopesticide on larvae of non-target *Drosophila* species present in *Bt*-treated areas. Doses up to those recommended for field application (10^6^ CFU/g of fly medium) had no effect on the fly development, whereas doses 10 to 100-fold higher (10^7^-10^8^ CFU/g) increased developmental time and decreased adult emergence rates in a dose-dependent manner and with varying effect amplitudes for all the species and strains tested. For all them, all larvae died before pupation at the highest dose tested (10^9^ CFU/g). Focusing on *D. melanogaster*, delayed development and reduced emergence resulted from stage-dependent larval mortality, and fitness-related traits of adult flies emerging from surviving *Bt* biopesticide exposure were moderately increased. The effects of *Bt* biopesticide seemed to result from the spores/cleaved toxins synergy, and possibly additives. While recommended doses had no effect on non-target *Drosophila* species, misuse or local accumulation of *Bt* bioinsecticides in the environment could have non-intentional side-effects on fly populations with potential implications for their associated communities.

## Introduction

The world’s population is expected to reach more than 9 billion people by 2050,^[1]^ increasing the demand for agricultural resources. This requires to improve pest management, especially insects that cause more than 30% of agricultural losses.^[2]^ Nowadays, their management largely relies on conventional chemical insecticides. However, their use and efficiency have been considerably reduced due to the emergence of pests’ resistance, development of secondary pests, adverse side-effects on non-target species (natural enemies of pests, pollinators),^[3,4]^ and more generally the impacts on biodiversity and human health (e.g. neurological disorders, functional impairment of reproduction, cancers).^[5-8]^ Developed as a more specific and safer alternative, biopesticides represent less than 5% of the pesticide market, the large majority being microbial insecticide formulations based on viable spores and toxins of the bacterium *Bacillus thuringiensis* (*Bt*) (over 400 registered formulations).^[4,9]^

*Bt* is a Gram-positive endospore-forming bacterium that synthesizes a wide range of toxins with different chemical structures, modes of action and biological targets. The most abundant and studied are Cry d-endotoxins encoded by genes located on large plasmids and produced as parasporal crystalline inclusions during the stationary growth phase.^[10,11]^ *Bt* produces other insecticidal toxins, the Cyt (cytolytic d-endotoxins) and Vip (secreted Vegetative Insecticidal Proteins) that synergize their effects with Cry toxins, virulence factors such as β-exotoxins (or thuringiensin), a secreted nucleotide toxic for almost all tested life forms thus prohibited in commercial formulations,^[12]^ and anti-fungal factors.^[13,14]^ *Bt* subspecies and strains can differ in their plasmid number and in the synthesized toxins cocktail responsible for their biological activity, which was used to delineate potential target insects.^[15]^ For instance, *Bt* subsp. *kurstaki* (*Btk*) produces the 5 Cry toxins, Cry1Aa, Cry1Ab, Cry1Ac, Cry2Aa and Cry2Ab,^[10,16]^ while *Bt* subsp. israelensis (*Bti*) produces a combination of Cry4Aa, Cry4Ba, Cry10Aa, and Cry11Aa,^[17,18]^ both strains being commercially used. The different toxin cocktails produced by some *Bt* subspecies can also be detrimental to non-insect organisms such as nematodes, protozoa, and even molluscs.^[15]^

The bioinsecticide formulations based on spores and toxin crystals of *Btk* and *Bti* are the most sprayed in organic and conventional farming, and natural areas (e.g. forests, swamps) to deal with larvae of Lepidopteran pests and Dipteran larvae of mosquitoes and black flies, respectively. It is generally accepted that once ingested by insect larvae, the toxin crystals are dissolved by the midgut alkaline pH, releasing ∼130 kDa pro-toxins that are then processed by digestive proteases into smaller, soluble, active toxin fragments of ∼ 60-70 kDa.^[19,20]^ Active toxins bind to specific receptors of midgut epithelial cells, eliciting pores formation in the cell membrane, cell lysis and gut epithelium disorganization.^[21,22]^ This allows gut bacteria, including *Bt*, to colonize the hemocoel, and leads to rapid septicaemia and death.^[23,24]^

Numerous impact studies of field application rates and acute intoxications have shown that *Bt* bioinsecticides are safe or have limited impact on non-target vertebrates and invertebrates, and associated species communities.^[25,26]^ However, the increasing use of biopesticides based on *Bt* spores and toxins has recently raised concern^[27]^ and led to the assessment of their potential effects on non-target species, such as auxiliary insects of biological control,^[28]^, pollinators,^[29]^ and species and species communities which simply share their habitat with *Bt*-targeted insect pests.^[30-32]^ Yet, there is growing evidence of direct and indirect cross-effects of *Bt* bioinsecticides and toxins across insect species and orders, or even across phyla, suggesting that *Bt* targeting is only partly specific.^[33,34]^ Data showed that almost all of the applied *Btk* dose was still present on the leaves surface 72 hours after spraying,^[35]^ its amount returning close to environmental levels only 28 days after treatment.^[36]^ Finally, *Bt* spores can survive in the soil and on different supports for months and even years after application.^[37-41]^ *Bt* formulations contain also numerous compounds to protect spores and crystals and aggregate them into a wettable form, surfactants to facilitate spraying and dispersion on plants, and phagostimulants.^[42,43]^ Nevertheless, since toxin crystals, and to a much lesser extent spores,^[44]^ are somewhat sensitive to abiotic conditions (e.g. UV, pH, rainfall), repeated sprayings with a minimum delay of 3 to 8 days is often recommended over the period of pest occurrence to achieve the required pest control level^[43,45]^ (http://www.certiseurope.fr, http://www.certisusa.com). All these can potentially lead to *Bt* accumulation in the environment, thus raising the rarely addressed issue of potential side-effects of chronic exposure (*i.e.* continuous and increasing exposure dose for an extended period) of non-target species to doses unexpectedly above the recommended spraying doses.

Diptera are worldwide distributed insects, most of which are not targets for *Bt* and its toxins. This is the case of the genus *Drosophila*, represented by ∼ 1 500 described species,^[46]^ including the model organism *D. melanogaster*. In the field, most of these flies feed and reproduce mainly on ripening or rotting/fermenting fruits and are therefore naturally present in areas treated with *Bt* such as orchards, vineyards and gardening areas. Unable to disperse between food patches, early developmental stages of *Drosophila* eat intensively and grow exponentially,^[47]^ and may thus ingest high doses of *Bt* bioinsecticides that have accumulated during the treatment periods. Surprisingly, despite the presence of many *Drosophila* species in *Bt-*treated areas, their role in the decomposition of organic matter, and the ease of study of some species, only a few studies have focused on these flies. However, most of them suggested susceptibility to *Btk*, but they used mainly late 3^rd^ instar larvae preparing for pupation, which do not feed much, and used *Bt* preparations, especially field isolates, that possibly contained highly toxic β-exotoxins, which are not authorized in commercial *Bt* formulations.^[48-55]^ So far, no study addressed the effects of chronic exposure to *Bt* formulations containing spores and toxin crystals but no β-exotoxins, on developing stages of these Dipterans that are present in *Bt*-treated areas.

Here, we have tested the chronic side-effects of different commercial formulations of *Btk* (devoid of β-exotoxins) and, to a lesser extent of *Bti*, with doses starting from recommended spraying doses up to 1 000 times this dose (i.e. below acute intoxication doses used in most studies). We mainly focused on developmental traits (developmental time, emergence rate), firstly using the wild-type *D. melanogaster* Canton S. The spore-forming Gram-positive *Bacillus subtilis* and the *Btk* strain (4D22), devoid of Cry toxin genes and thus of crystals, were used as non-pathogenic controls. We also analysed two fitness-related traits of adult flies (male and female longevity, offspring number) after entire development in presence of *Btk* formulation. To test for effects specific to the fly genetic background, developmental traits upon exposure to *Btk* formulation were measured on four other *D. melanogaster* strains. Finally, we extended further these development experiments to seven other *Drosophila* species. We have chosen six species in addition to *D. melanogaster*, including cosmopolitan species, which can co-occur in the field^[56-60]^ and are usually present in *Bt*-treated areas, and the invasive *D. suzukii* that can now co-exist with the six species in the areas it has recently invaded. This aims at providing a first-step in the exploration of potential implications of chronic exposure to *Btk* formulation in terms of species competition and species community composition and dynamics.

## Material and methods

### Commercial formulations, Bacillus productions and Colony Forming Unit measurement

The tested commercial brands of *Bacillus thuringiensis kurstaki* (*Btk*; serotype 3a, b, c^[61]^) were Delfin^®^ A and B (strain SA-11; wettable granules, Valent BioSciences, AMM 9200482, 32,000 UI/mg) and Scutello DF (a Dipel^®^ sub-brand; strain ABTS-351; wettable granules, Biobest^®^, AMM 2010513, 540g/kg). The commercial brand of *Bacillus thuringiensis israelensis* (*Bti*; strain HD-14; serotype 14^[61]^) was VectoBac^®^ WG (wettable granules, Bayer, AMM 2020029, 3000 UTI/mg). For each formulation, the number of viable spores (expressed as Colony Forming Units (CFU) per mg of granules) was estimated using serial dilutions of a suspension on LB agar plates and counting of bacterial colonies after overnight incubation at 30°C. CFU estimations were 5×10^7^ CFU/mg for *Btk* Delfin^®^ A; 2.5×10^7^ CFU/mg for *Btk* Delfin^®^ B; 2.2×10^7^ CFU/mg for *Btk* Scutello DF; 6×10^7^ CFU/mg for *Bti* VectoBac^®^. Of note, our CFU estimations fell in those appended on the commercial packaging, between 10^13^ and 5×10^13^ CFU/kg (32 000 UI/mg; http://www.certiseurope.fr). No change in CFU estimations occurred during the time frame of the experiments. Manufacturer-recommended doses for Delfin^®^ range from 0.15 to 1.5 kg/ha depending on the crop type. Based on our CFU estimations, this corresponds to recommended doses of 7.5×10^4^ to 7.5×10^5^ CFU/cm^2^ of Delfin^®^ A, and 3.75×10^4^ to 3.75×10^5^ CFU/cm^2^ of Delfin^®^ B for each spraying in the field. For Scutello DF, recommended doses range from 0.1 to 1 kg/ha, which are equivalent to 2.2×10^4^ to 2.2×10^5^ CFU/cm^2^. Vectobac^®^ WG is used at 0.125 to 1 kg/ha, equivalent to 7.5×10^4^ to 6×10^5^ CFU/cm^2^.

The acrystillipherous (Cry toxin-free) *Btk* 4D22 strain (depleted for the toxin-encoding plasmids^[62]^) obtained from the Bacillus Genetic Stock Center (http://bgsc.org; Columbus USA), and a *Drosophila* non-pathogenic *Bacillus subtilis* (from Dr. E. Bremer, University of Marburg, Germany) were grown at 30°C in the sporulation-specific medium PGSM (Bactopeptone^®^ 7.5 g, KH_2_PO_4_ 3.4 g, K_2_HPO_4_ 4.35 g, glucose 7.5 g, PGSM salts 5 mL, CaCl2 0.25 M, distilled water qsp 1L, pH 7.2; PGSM salts: MgSO_4_.7H_2_O, MnSO_4_.H_2_O, ZnSO_4_.7H_2_O, FeSO_4_.7H_2_O) for about 14 days for sporulation to occur. Following elimination of vegetative cells (1h at 70 °C), spore pellets were collected after centrifugation (4,500 rpm, 20 min, 4 °C), washed with sterile water, and lyophilized. CFU numbers were counted for each preparation as described above.

### Fly stocks

The four tested strains of *Drosophila melanogaster* (phylogenetic subgroup: melanogaster) were the standard wild-type Canton S (Bloomington Drosophila Centre) used as a reference strain, the wild-type Nasrallah strain from Tunisia (strain 1333, Gif-sur-Yvette), the double mutant standard strain YW1118 (white and yellow mutations; gift from Dr. B. Charroux, IBD, Marseille-Luminy), and a recently field-collected strain (caught in Southern France in 2013) that we named “Sefra”. For *Drosophila* species comparison, we included 6 species of the *Drosophila* subgenus, *D. simulans* (strain 1132; phylogenetic subgroup: melanogaster), *D. yakuba* (strain 1880; phylogenetic subgroup: melanogaster), *D. hydei* (phylogenetic subgroup: hydei) and *D. suzukii* (phylogenetic subgroup: immigrans) (both kindly provided by Dr. R. Allemand, LBBE, University Lyon 1), *D. immigrans* (phylogenetic subgroup: immigrans), *D. subobscura* (phylogenetic subgroup: obscura), and one species of the *Dorsilopha* subgenus, *D. busckii* (all three species collected in South-East of France in Spring 2015).

All strains and species were maintained at controlled densities (150-200 eggs/40 ml of fly medium) under standard laboratory conditions (25°C or 20°C for recently collected species, 60 % relative humidity, 12:12 light/dark cycle), on a high-protein/sugar-free fly medium (10 % cornmeal, 10 % yeast, 0 % sugar). The *D. melanogaster* Canton S strain was also reared on a standard low-protein/sugar-free fly medium (8 % cornmeal, 2 % yeast, 2.5 % sugar) to test for the influence of the medium composition on *Btk* exposure effects.

### Intoxication method and dose-response assay

Commercial formulations and laboratory spore productions were suspended and diluted in buffer to perform dose-response assays with doses from 10^5^ to 10^9^ CFU/g of fly medium. All doses were prepared in 100 µl and homogenized thoroughly with the fly medium (100µl/g). *Drosophila* eggs and larvae were collected from stock vials at the suitable developmental stage and transferred carefully to the intoxication vials and dishes, then maintained under standard laboratory conditions until a) the emergence of adults, or, in the larvae survival tests, b) until a given developmental stage was reached from the egg, and c) for 24h. Control groups of individuals were transferred on fly medium homogenized with the same volume of buffer.

### Development-related traits and larval survival

To evaluate emergence rates and developmental times upon intoxication throughout the entire development, precise numbers of eggs from mass oviposition were transferred to intoxication vials containing fly medium mixed with doses of *Bt* formulations or bacteria productions and let to develop under standard laboratory conditions until the fly emergence. Eggs without chorion and transparent eggs were discarded. The initial number of eggs was adjusted depending on the species biology and the vial size: 20 eggs for 2 g of fly medium in small vials (Ø 3.3 cm, surface ∼8.5 cm^2^, 0.24 g/cm^2^) for tests with *D. melanogaster* Canton S, 50 eggs for 6 g of fly medium for comparison of *D. melanogaster* strains and *Drosophila* species in wider vials (Ø 4.6 cm, surface ∼16 cm^2^, 0.37 g/cm^2^) except for *D. hydei, D. suzukii* and *D. immigrans* for which 30 eggs were transferred on 6 g of fly medium. Numbers and sex of emerging flies were recorded once a day until the day the pupae of the next generation should form. From these data, the emergence rate (proportion of emerged flies from the initial eggs; ER), the mean developmental time (mean number of days for completion of development; DT), and the sex-ratio (proportion of male flies; SR) were calculated for each intoxication vial.

For the larval survival tests, 20 eggs or larvae from a 4-hour mass oviposition at the indicated developmental stage, were transferred to small dishes containing 1 g of fly medium (Ø 3 cm, surface ∼7 cm^2^) homogenized with increasing doses of Delfin^®^ A. Surviving larvae were counted at the indicated developmental stage, or after 24-hour intoxication, to calculate the proportion of surviving larvae. For the test from the egg, eggs which did not hatch were not included in the counting. As a control measurement, we measured the pH of the fly medium in the presence of the dose range of *Bt* formulations (see Supplementary Information S4).

### Adult fitness-related traits

For the longevity and offspring number tests, males and females emerged from several rearing vials for each dose of Delfin^®^ A were pooled when aged 2 days. Groups of 15 males and 15 females were transferred into vials with fresh fly medium without formulation. Fly medium was renewed every 3-4 days. After each fly transfer to fresh food, discarded maintenance vials were incubated under standard laboratory conditions for the offspring to develop. Mortality and sex of dead flies were recorded daily until the last fly died. Offspring numbers were counted from the first emergence until pupae of the next generation appeared. The tests were repeated twice. Due to the variation in the duration of the two longevity experiments, offspring numbers of each vial were summed to obtain a total offspring number per dose of Delfin^®^ A for each experiment.

### Dialysis and Cry toxin analysis

A suspension of 2×10^10^ CFU of Delfin^®^ A was dialyzed against PBS (KH_2_PO_4_ 1.06 mM, Na_2_HPO_4_(2H_2_O) 3mM, NaCl 154 mM, qsp distilled water, pH 7.2), at 250 rpm, 4°C overnight, using an 8-10 kDa MW cut-off membrane (ZelluTrans, Roth^®^). The CFUs of the dialyzed suspension and the effects on emergence rate (ER) and developmental time (DT) were analysed as described above. The dialyzed suspension was also subject to a 12.5 % SDS-PAGE and compared to the non-dialyzed suspension after silver staining. The presence of Cry1A pro-toxins, activated toxins and toxin fragments was probed by Western-blot using an in-house anti-Cry1A rabbit polyclonal antibody.

### Data analysis

Data on development traits (emergence rate (ER) and developmental time (DT)), sex-ratio (SR), survival of larval stages and offspring number were analysed with mixed effect models including the dose of *Btk* formulation/spore production, the *D. melanogaster* strain, the *Drosophila* species or the developmental stage as fixed effects, and replicate (plus the experiment when necessary) as random effects (for ER data, data were analysed with bias-corrected models with replicate as fixed effect to allow pairwise comparisons; similar results obtained with models including replicate as random effect). ER, SR and survival of larval stages were analysed with generalized linear models, with binomial distribution and logit link. DT and offspring number were analysed with linear models. DT were transformed into developmental rates (1/developmental time) to fulfil the assumptions of the analysis of variance (homoscedasticity and residuals normality). Adult longevity data were analysed with proportional hazard Cox regression models including fly sex and dose of *Btk* formulation as fixed effects, and replicates as a random effect. For all the data sets, the main fixed effects and their interactions were tested with log-likelihood ratio tests. *Post hoc* pairwise comparisons were made for pairs of *D. melanogaster* strains, formulation/spore treatments, and between the control dose and the other doses. All the analyses were performed in R^[63]^ using the packages lme4,^[64]^ brglm,^[65]^ multcomp,^[66]^, survival,^[67]^ and coxme.^[68]^

## Results

### *Btk* formulations adversely impact the development of *D. melanogaster*

The *D. melanogaster* wild-type Canton S strain was used to evaluate the dose-dependent effect of the commercial *Btk* formulation Delfin^®^ A on the emergence rate (ER, proportion of emerged flies from the initial egg pool) and developmental time (DT, mean number of days from egg to adult emergence). Eggs were transferred on a standard low-protein/high-sugar fly medium containing Delfin^®^ A at doses ranging from 5×10^5^ CFU/g of medium (mean equivalent of the maximum recommended doses for one field application; see Methods and Supplementary information S1) to 10^9^ CFU/g (∼ 1,000 times the recommended dose). To check for specific effects of *Btk* formulations and the respective role of *Btk* spores and Cry toxins, we tested the same dose range of the commercial *Bti* formulation Vectobac^®^ targeting mosquitoes that contains different Cry toxins,^[22]^ of the Cry-free strain *Btk* 4D22, and of the *Drosophila* non-pathogenic spore-forming *Bacillus subtilis*.

Developmental traits (ER and DT) of exposed and non-exposed control flies were similar at doses up to 10^7^ CFU/g of Delfin^®^ A (Fig. 1a-b; Table 1). At higher doses, both ER and DT were affected in a dose-dependent manner: ER was reduced by 17% at 5×10^7^ CFU/g (although not statistically significant), up to 100% at 10^9^ CFU/g, dose at which no individual reached the pupal stage. The lethal dose 50 (LD50) was estimated between 5×10^7^ and 10^8^ CFU/g (Fig. 1a). DT was increased of about 0.5 day at 5×10^7^ CFU/g (+ 4% compared to controls), up to 1.5 days (+ 14%) at 10^8^ CFU/g (Fig. 1b; Table 1). The sex-ratio at emergence (SR, proportion of males) was strongly biased towards males at the highest dose at which complete development occurred (10^8^ CFU/g), with 58% more males compared to controls (Supplementary information S2). Because addition of *Btk* formulation could modify parameters of the fly medium and thus contribute to these effects, we checked the pH of the medium: the presence of formulation and its dose had no effect on it (Supplementary information S4).

**Table 1.**
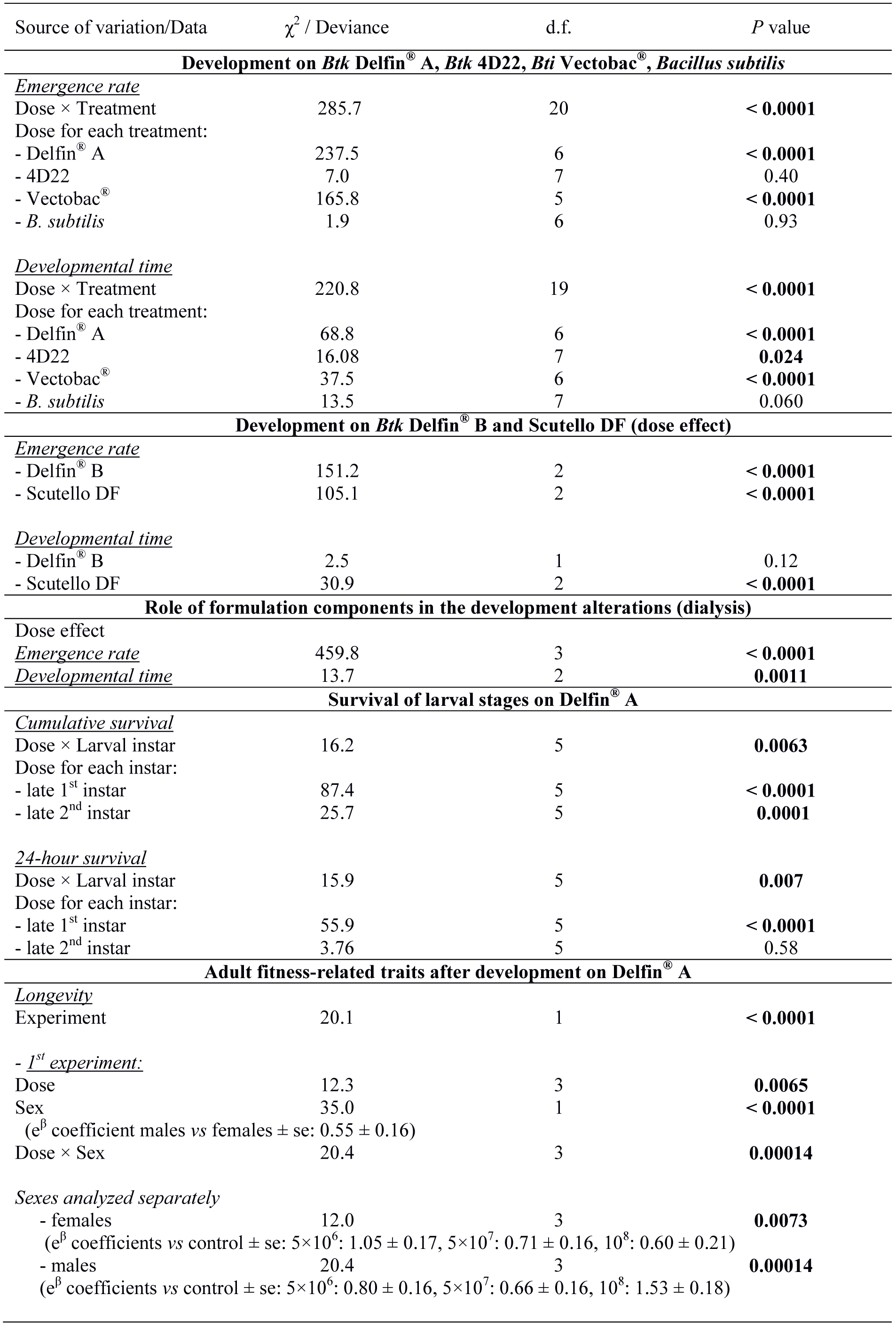
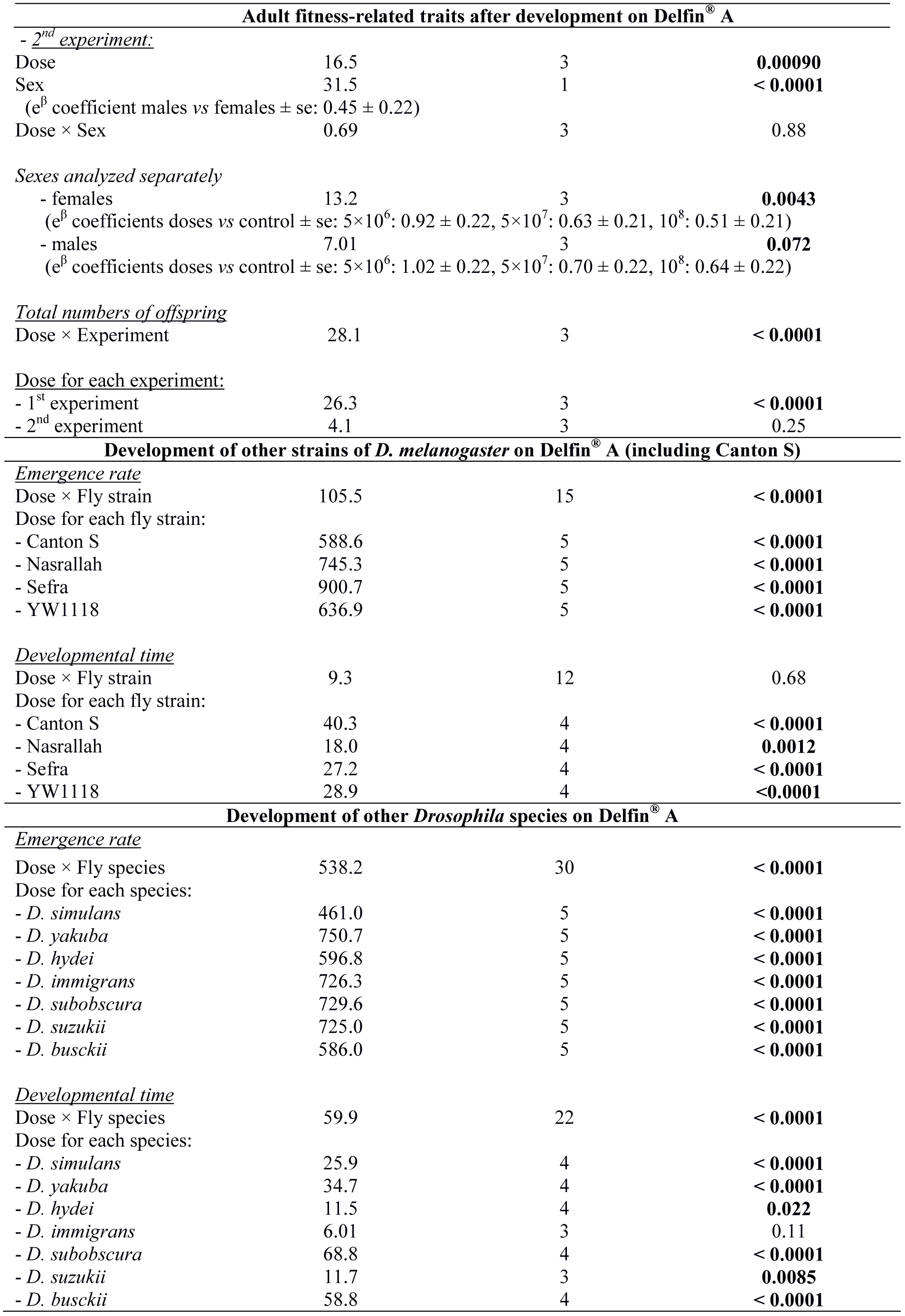
Results of statistical analyses to assess the effect of the dose of formulation/spore production and its interaction with the treatment, the larval instar, the experiment, the sex, the fly strain and the fly species when appropriate. See figures for *post hoc* comparisons of the doses with the control dose.

**Figure 1.**
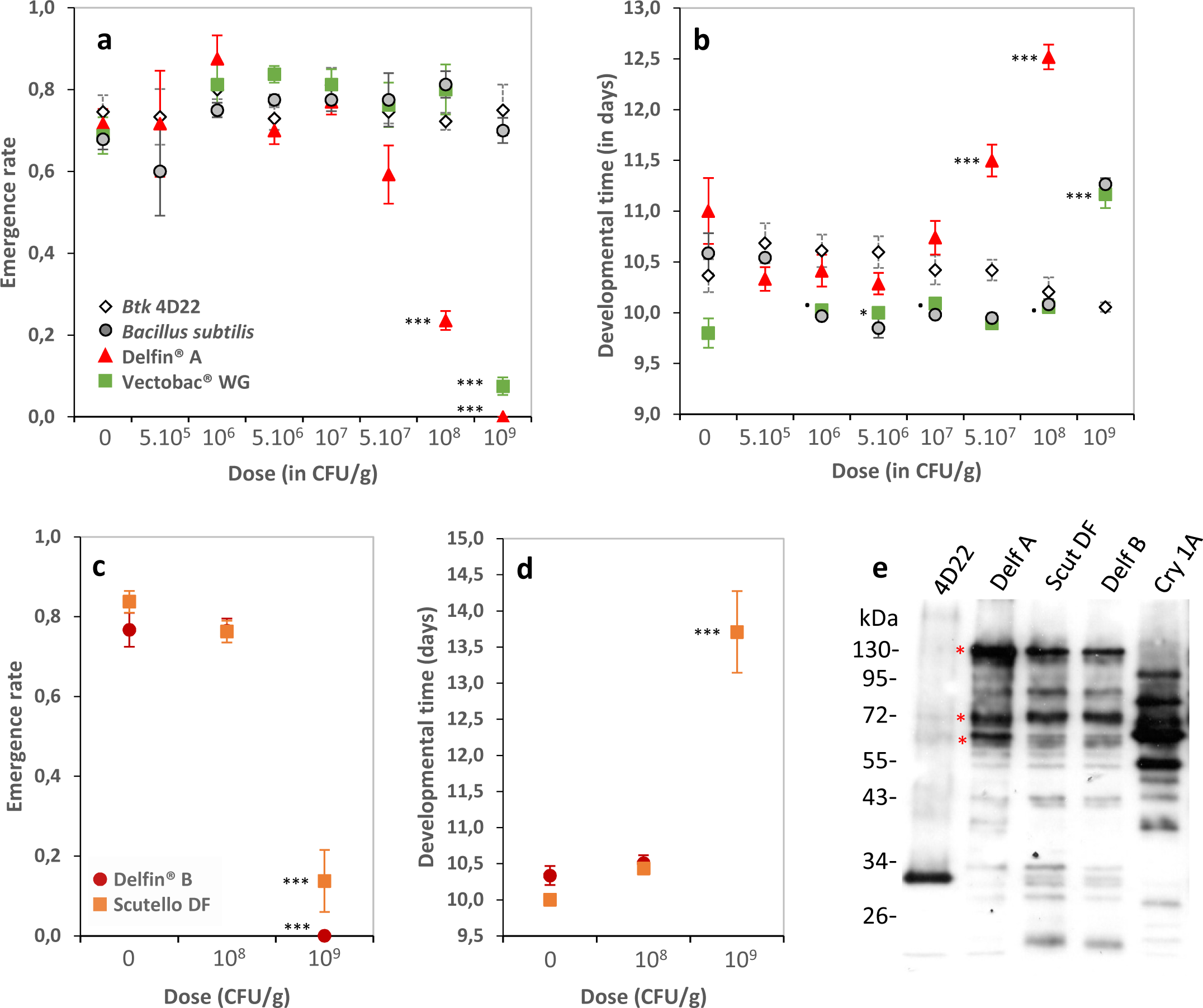
Development of *D. melanogaster* Canton S flies on *Btk* and *Bti* commercial formulations. (**a**) Emergence rate (mean ± s.e.m.) and (**b**) developmental time (mean ± s.e.m.) of 20 initial eggs on increasing doses of *Btk* Delfin^®^ A (red triangles), the Cry-free *Btk* 4D22 (open lozenges), the mosquito-targeting *Bti* Vectobac^®^ (green squares), and the non-pathogenic *Bacillus subtilis* (light grey circles). For Vectobac^®^ and *B. subtilis, N* = 4-7 per dose; for Delfin^®^ A and *Btk* 4D22, *N* = 9-12 for the control, *N* = 3 for 5.10^5^ and 10^9^, *N* = 4-9 for 10^6^, *N* = 7-14 from 5.10^6^ to 10^8^. (**c**) Emergence rate (mean ± s.e.m.) and (**d**) developmental time (mean ± s.e.m.) on increasing doses of the two *Btk* formulations Delfin^®^ B (dark red circles) and Scutello DF (orange squares). *N* = 4 replicates of 20 eggs per dose and formulation, except for controls and 10^8^ CFU/g of Delfin^®^ B (9-10 replicates of 20 eggs). Results of *post hoc* comparisons of each dose to the control: · 0.05<*P*<0.1; * 0.01<*P*<0.05; ** 0.001<*P*<0.01; *** *P*<0.001. (**e**) Immunoblotting with an anti-Cry1A polyclonal antibody on proteins from a suspension of laboratory-produced spores of Cry-free *Btk* 4D22, the three *Btk* formulations Delfin^®^ A, B, Scutello DF, and a suspension of laboratory-produced Cry1A toxins. Red asterisks indicate the Cry protoxins (∼130 kDa) and the activated fragments (∼60 kDa and ∼70 kDa).

We observed no change in ER using the same dose range of the *Btk* Cry-free strain 4D22 (Fig. 1a, 1e; Table 1) and the non-pathogenic *Bacillus subtilis* (Fig. 1a, Table 1). Addition of *Bti* Vectobac^®^ did not affect ER up to 10^8^ CFU/g but reduced it by 89% at 10^9^ CFU/g (∼2,000 times the highest recommended dose for field application; Fig. 1a; Table 1; Supplementary information S1). DT varied with the dose of *Btk* 4D22, the differences being mainly between doses but not with the control. DT increased by ∼1.5 days at the highest dose of Vectobac^®^ (Fig. 1b; Table 1) and showed a similar trend with *B. subtilis* (*P* = 0.06; Fig. 1b; Table 1). None of these three treatments influenced dramatically the SR, the slight decrease in male proportion for most of the Vectobac^®^ doses being due to the higher average sex-ratio for the control dose compared to those for the two other treatments (Supplementary information S2).

To test whether these effects are generic to *Btk* formulations, the fly development was evaluated on two other formulations, Delfin^®^ B (same brand) and Scutello DF (brand Dipel^®^), at the critical doses 10^8^ and 10^9^ CFU/g. As Delfin^®^ A, these formulations contain spores and Cry toxins such as Cry-1A as pro-toxins of ∼130 kDa, activated toxins of ∼60-70 kDa, but also as smaller fragments^[20]^ (Fig. 1e, red asterisks). ER remained unchanged at 10^8^ CFU/g whereas no individual reached pupation at 10^9^ CFU/g on Delfin^®^ B and very few individuals reached the adult stage on Scutello DF^®^, DT being increased by more than 2 days (Fig. 1c-d; Table 1). No significant bias in SR was observed for either formulation (Supplementary information S2).

### Adverse effects of *Btk* formulation strongly impact the early development

Larval stages were assessed for their susceptibility to *Btk* formulation in two independent and complementary dose-response tests of survival to Delfin^®^ A, at doses ranging from 10^5^ to 10^9^ CFU/g of high protein/sugar free medium (this medium is used to rear fly species which are difficult to rear in the lab (see below) and is less limiting for the development of early larval stages). We focused on the 1^st^ and 2^nd^ larval instars, during which growth is exponential,^[47]^ so that larvae are most heavily exposed to the bioinsecticide. In the first test, the cumulative survival was measured by counting alive late 1st and 2nd instar larvae which have been exposed to Delfin^®^ A from the egg stage. Larval survival was not influenced at 10^7^ CFU/g, whereas it decreased for both larval instars above that dose to reach up to 37% mortality at 10^9^ CFU/g (Fig. 2a). Reduced survival tended to occur at a lower dose when cumulative survival was measured later in the development, *i.e.* 10^9^ for late 1^st^ instar larvae and 10^8^ CFU/g for 2^nd^ instar larvae (Fig. 2a; Table 1). For both instars, larvae surviving 10^9^ CFU/g were noticeably smaller and less active than those surviving lower doses. In emergence assays with planned exposure from the egg to the adult stage, none of these individuals reached the pupal stage (see results above). In the second test, larval survival was measured after early 1^st^ and 2^nd^ instar larvae had been exposed for 24 hours to Delfin^®^ A. Survival of 1^st^ instar larvae decreased by 36% on 10^9^ CFU/g whereas that of 2^nd^ instar larvae did not change (Fig. 2b, Table 1).

**Figure 2.**
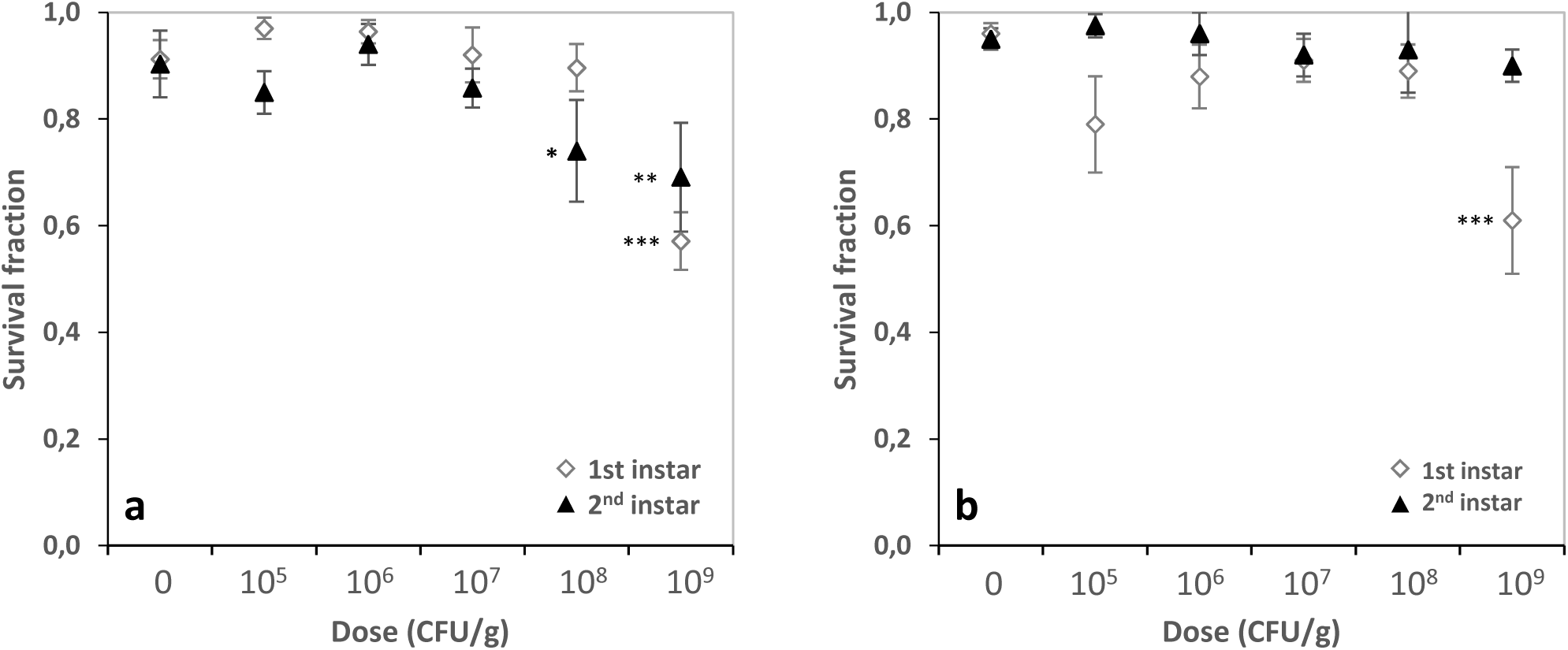
Survival of *D. melanogaster* Canton S larval stages on increasing doses of *Btk* Delfin^®^. **A.** (**a**) Proportion of surviving larvae (mean ± s.e.m.) upon *Btk* exposure from the egg to late 1^st^ instar (open lozenges) and late 2^nd^ instar (black triangles). (**b**) Proportion of surviving larvae (mean ± s.e.m.) upon 24-hour *Btk* exposure of early 1^st^ instar larvae (open lozenges) and 2^nd^ instar larvae (black triangles). *N* = 5-7 replicates of 20 individuals per dose. Results of *post hoc* comparisons of each dose with the control: * 0.01<*P*<0.05; ** 0.001<*P*<0.01; *** *P*<0.001.

### Developmental exposure to *Btk* formulation does not strongly influence fitness-related traits in adults

Long-term consequences on flies of exposure to *Btk* formulation throughout the development were evaluated on two fitness-related traits, longevity and total offspring number. Traits were measured on a *Btk*-free low-protein/high-sugar medium after individuals had completed their development on the same fly medium but in presence of selected doses of Delfin^®^ A: 5×10^6^ CFU/g, which had no impact on development, and 5×10^7^ and 10^8^ CFU/g, which caused moderate and strong development alterations, respectively (see Fig. 1a).

Adult longevity was analysed in two independent sets of experimental replicates on groups of 15 females and 15 males held together. Despite large variation between the two sets of experimental replicates (Table 1), the longevity of adults reared on 5×10^6^ CFU/g of Delfin^®^ A was similar to that of non-exposed controls (Fig. 3). Males and females which developed on the two higher doses showed a moderate longevity benefit, higher in females for 10^8^ CFU/g (Fig. 3a-b, d-e; Table 1). Males generally survived better than females (Table 1) but their longevity benefit of developing on 10^8^ CFU/g was only observed in the second experiment (Fig. 3b, e).

**Figure 3.**
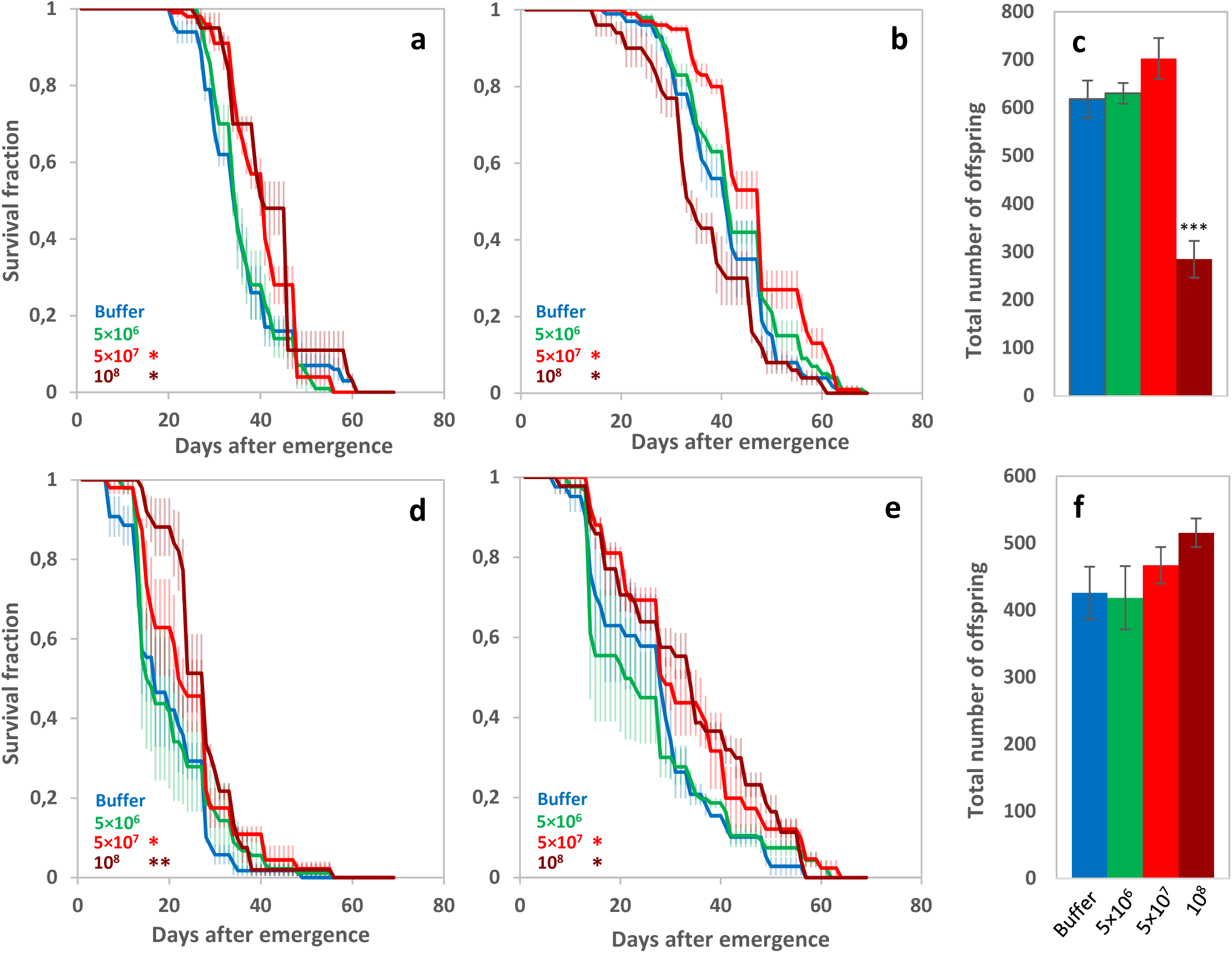
Fitness-related traits of adults (longevity and total offspring number) after development on *Btk* Delfin^®^. **A.** (**a, d**) Female longevity (mean survival fraction over time ± s.e.m.), (**b, e**) Male longevity (mean ± s.e.m.), and (**c, f**) total offspring number (mean ± s.e.m.), measured on individuals that developed without *Btk* (blue items) and on 5×10^6^ CFU/g of *Btk* Delfin^®^ A (green items), 5×10^7^ CFU/g (red items), and 10^8^ CFU/g (dark red items). Data from 2 experiments (a-c, experiment 1; d-f, experiment 2). For each trait, *N* = 3-5 replicates of 15 males and 15 females per dose in experiment 1, *N* = 3 replicates of 15 males and 15 females in experiment 2. Results of *post hoc* comparisons of each dose with the control: * 0.01<*P*<0.05; ** 0.001<*P*<0.01; *** *P*<0.001.

The female offspring number - the sum of offspring produced by the 15 females of each fly group during the longevity experiment - varied depending on both the experiment and the Delfin^®^ A dose (Table 1). In the 1^st^ experiment, adults from larvae reared on 10^8^ CFU/g had fewer offspring compared to control adults and to adults developed on the other doses whereas the total offspring number varied regardless of the *Btk* dose in the 2^nd^ experiment (Fig. 3c, f, Table 1).

### *Btk*-formulation dose-dependent alterations of development are not specific to the *D. melanogaster* Canton S strain

Dose-dependent effects of *Btk* formulation on the development were tested on three additional *D. melanogaster* strains: the wild-type Nasrallah (strain 1333), the wild-type Sefra population reared in the laboratory for 4 years, and the double mutant YW1118. The emergence rates (ER) and developmental times (DT) were measured on a high-protein/sugar-free medium (rearing medium of these strains) mixed with Delfin^®^ A doses ranging from 10^5^ to 10^9^ CFU/g. To allow the comparison with previous results with Canton S flies on low-protein/high sugar fly medium, Canton S was also reared and tested on the high-protein/sugar-free medium along with the other strains.

None of the fly strains was impacted at doses up to 10^7^ CFU/g, whereas ER was strongly reduced and DT was increased at higher doses for all the strains (Fig. 4a-b, Table 1), with no individual reaching the pupal stage at 10^9^ CFU/g (LD50 between 10^8^ and 10^9^ CFU/g). At 10^8^ CFU/g, the magnitude of effects on Canton S flies was lower than that observed on the low-protein/high-sugar medium. At this dose, ER varied between strains, the largest reduction being observed for Sefra (Table 1). We observed no dose-dependent bias in SR (Supplementary information S3).

**Figure 4.**
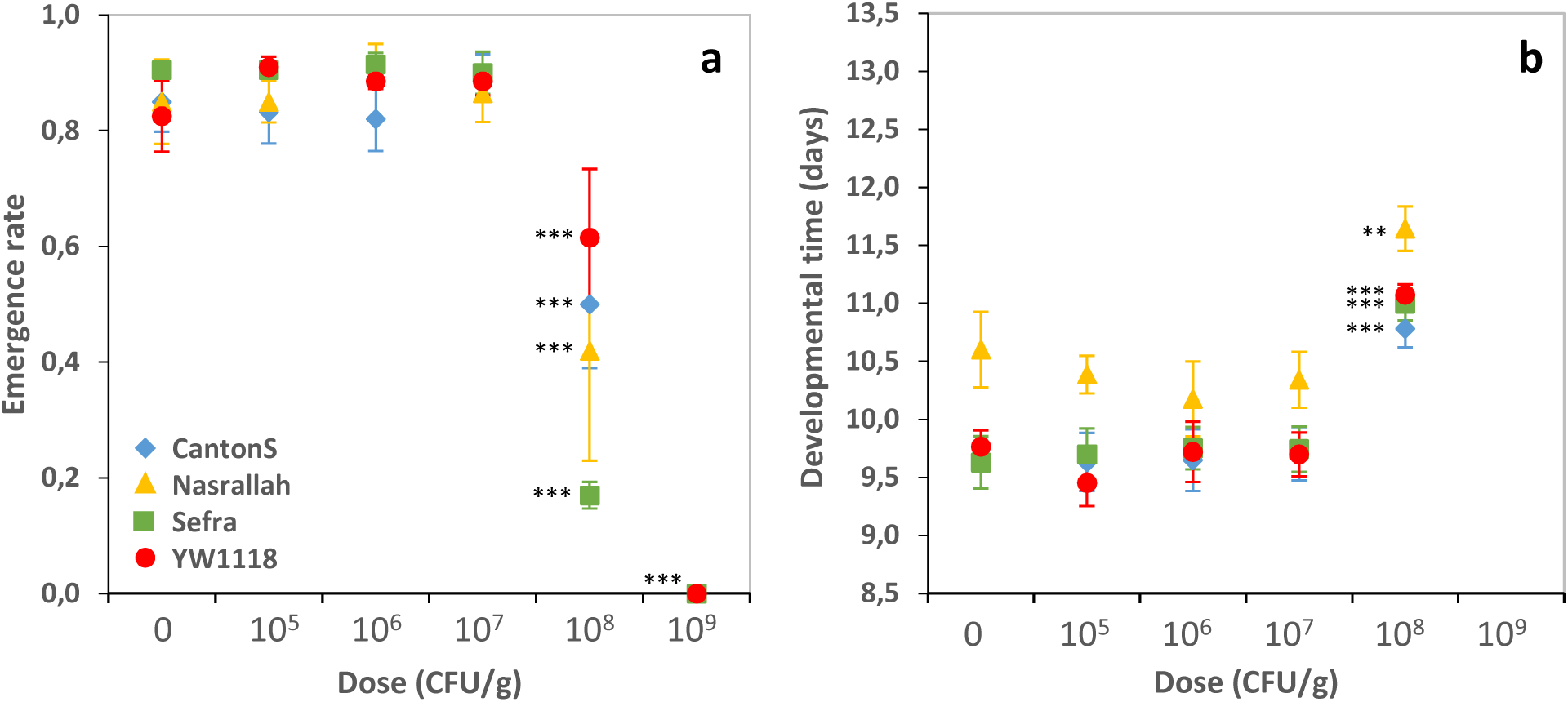
Development of four *D. melanogaster* strains on increasing doses of *Btk* Delfin^®^. **A.** (**a**) Emergence rate (mean ± s.e.m.), (**b**) Developmental time (mean ± s.e.m.) of the strains Canton S (blue lozenges), Nasrallah (yellow triangles), Sefra (green squares), and YW1118 (red circles). *N* = 4 groups of 50 eggs per dose and fly strain for each trait. Results of *post hoc* comparisons of each dose to the control: · 0.05<*P*<0.1; * 0.01<*P*<0.05; ** 0.001<*P*<0.01; *** *P*<0.001.

### *Btk* formulation affects differently other *Drosophila* species

The ER and DT were analysed for seven other *Drosophila* species from different phylogenetic clades at doses of Delfin^®^ A from 10^5^ to 10^9^ CFU/g of high-protein/sugar-free medium (rearing medium of all the species). Tested species were *D. simulans* (*D. melanogaster* sister species), the African *D. yakuba, D. subobscura, D. immigrans, D. hydei*, and the invasive *D. suzukii*, all belonging to the *Drosophila* subgenus, and *D. busckii* from the *Dorsilopha* subgenus. For all the species, doses up to 10^6^ CFU/g of Delfin^®^ A had no effect on ER and DT whereas all individuals failed to reach the pupal stage and no fly emerged at 10^9^ CFU/g (Fig. 5-6). Amplitudes of development alterations at 10^7^ and 10^8^ CFU/g varied between species (Fig. 5-6; Table 1). All species were affected at 10^8^ CFU/g as was *D. melanogaster* (see Fig. 4a for comparison). *D. simulans* and *D. busckii* had unchanged ER, but DT was slightly increased for *D. simulans* (although slightly reduced at 10^7^ CFU/g; similar results with a Japanese strain, data not shown) and strongly increased for *D. busckii* (by 20%, *i.e.* ∼ 4 days) (Fig. 5-6, Table 1). *D. yakuba* ER and DT were similar to those of *D. melanogaster*, with an LD50 around 10^8^ CFU/g and a moderate DT increase of ∼ 1 day (Fig. 5-6, Table 1; similar results with a strain from Sweden, data not shown). The ER of *D. hydei* and *D. subobscura* were very low at 10^8^ CFU/g (LD50 below this dose), with a high DT (Fig. 5-6; Table 1), while *D. immigrans* did not survive. No *D. suzukii* individual emerged at 10^8^ CFU/g and development was already moderately impacted at 10^7^ CFU/g (Fig. 5-6). No dose-dependent bias in SR was detected for either species (Supplementary information S5).

**Figure 5.**
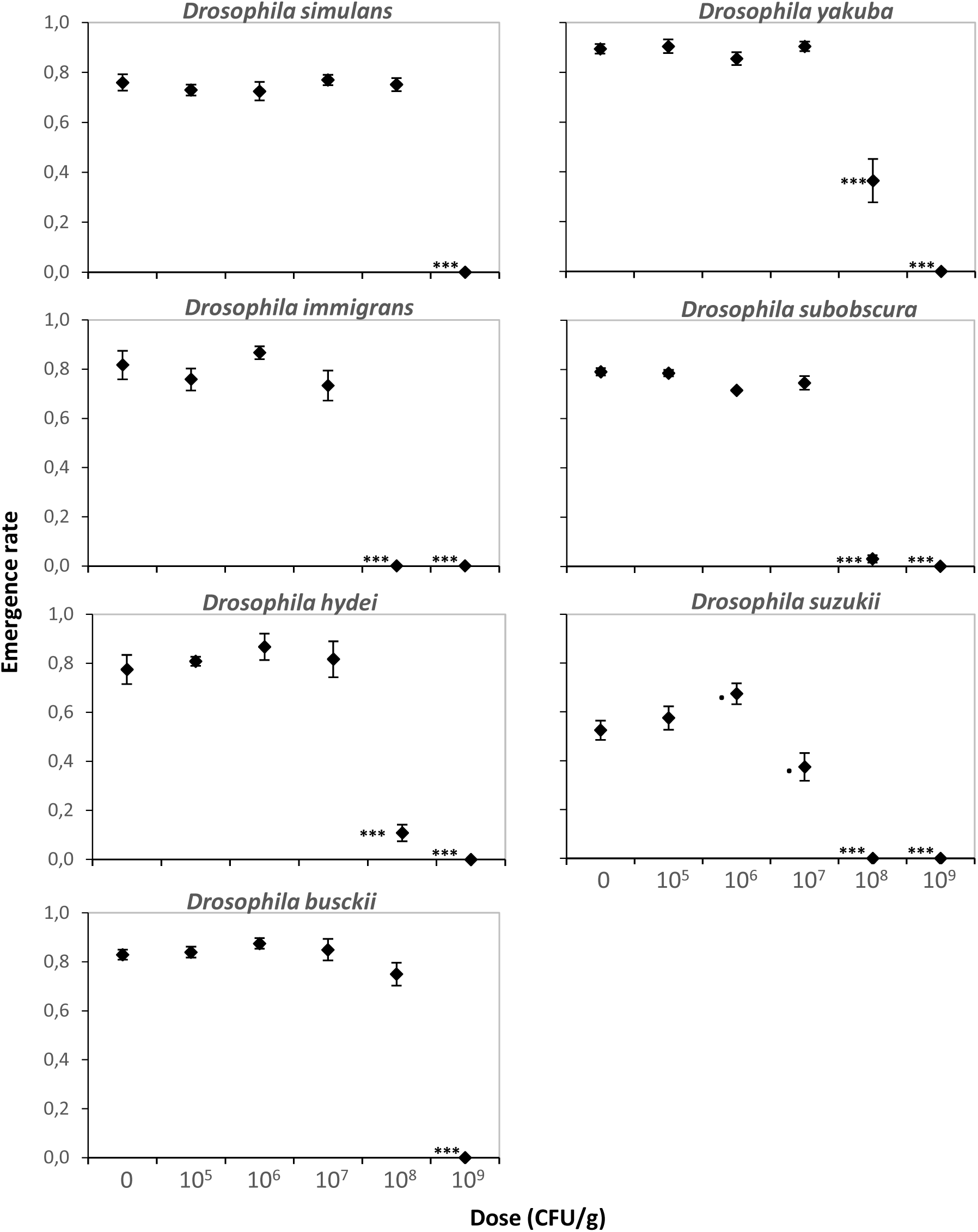
Emergence rate of seven *Drosophila* species on increasing doses of *Btk* Delfin^®^. **A.** Mean emergence rate (± s.e.m.). *N* = 4 replicates of 50 eggs per dose for *D. simulans, D. yakuba, D. subobscura*, and *D. busckii, N* = 4 replicates of 30 eggs per dose for *D. hydei, D. suzukii*, and *D. immigrans*. Results of *post hoc* comparisons of each dose with the control: · 0.05<*P*<0.1; * 0.01<*P*<0.05; ** 0.001<*P*<0.01; *** *P*<0.001.

### Development alterations may result from a synergy between formulation components

*Bt* spores and toxins can represent more than half the weight of commercial formulations (85% for Delfin^®^; http://www.certisusa.com), with up to about 10% of insecticidal protein toxins within this fraction, mainly Cry pro-toxins and activated toxins^[69]^ (see Fig. 1e). The remaining weight consists of various compounds such as residues of culture medium and various additives including surfactant, anti-foaming agents, etc…^[25,43]^ Since it has been shown that, for some products, additives can be more harmful in some cases than the active ingredient,^[70]^ we explored the role of small diffusible molecular weight components of Delfin^®^ A in the alterations of ER and DT of *D. melanogaster* Canton S. For that, we mixed a 10 kDa dialyzed suspension of Delfin^®^ A at 10^7^, 10^8^, and 10^9^ CFU/g with low-protein/high-sugar medium. ER and DT were unaffected by the presence of the dialyzed suspension at the 10^7^ CFU/g dose, whereas no individual reached the adult stage (no pupation) with the suspension at the 10^9^ CFU/g dose (Fig. 7a; Table 1). At 10^8^ CFU/g, ER was not modified but DT increased by ∼ 1 day, only in one of the two sets of experiments, partially reproducing the changes observed without dialysis (Fig. 7a-b; see also Fig. 1a-b, Table 1; 3 independent experiments for ER, 2 independent experiments for DT).

**Figure 6.**
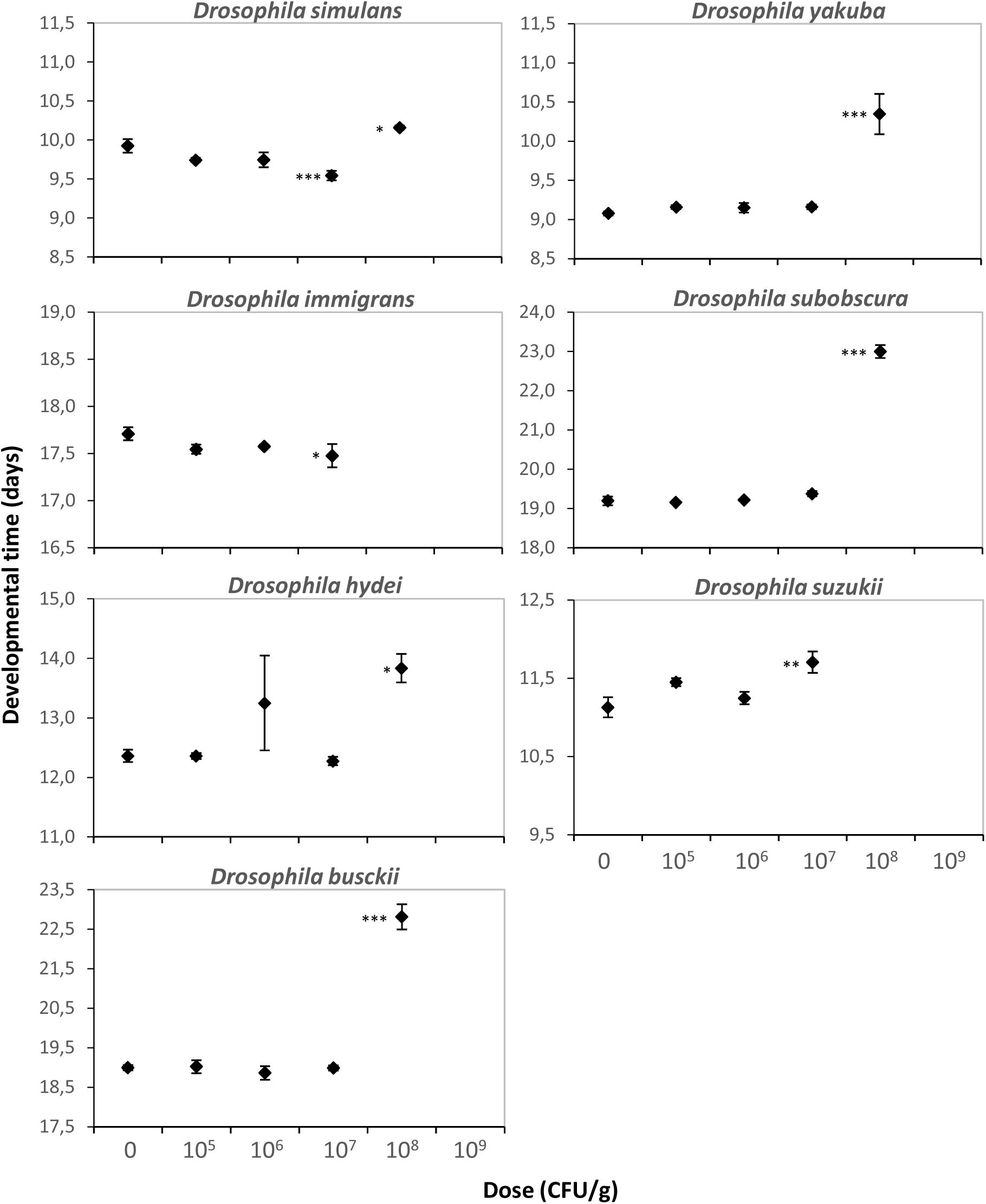
Developmental time of seven *Drosophila* species on increasing doses of *Btk* Delfin^®^. **A.** Mean developmental time (± s.e.m.). *N* = 4 replicates of 50 eggs per dose for *D. simulans, D. yakuba, D. subobscura*, and *D. busckii, N* = 4 replicates of 30 eggs per dose for *D. hydei, D. suzukii*, and *D. immigrans*. Results of *post hoc* comparisons of each dose with the control: * 0.01<*P*<0.05; ** 0.001<*P*<0.01; *** *P*<0.001.

**Figure 7.**
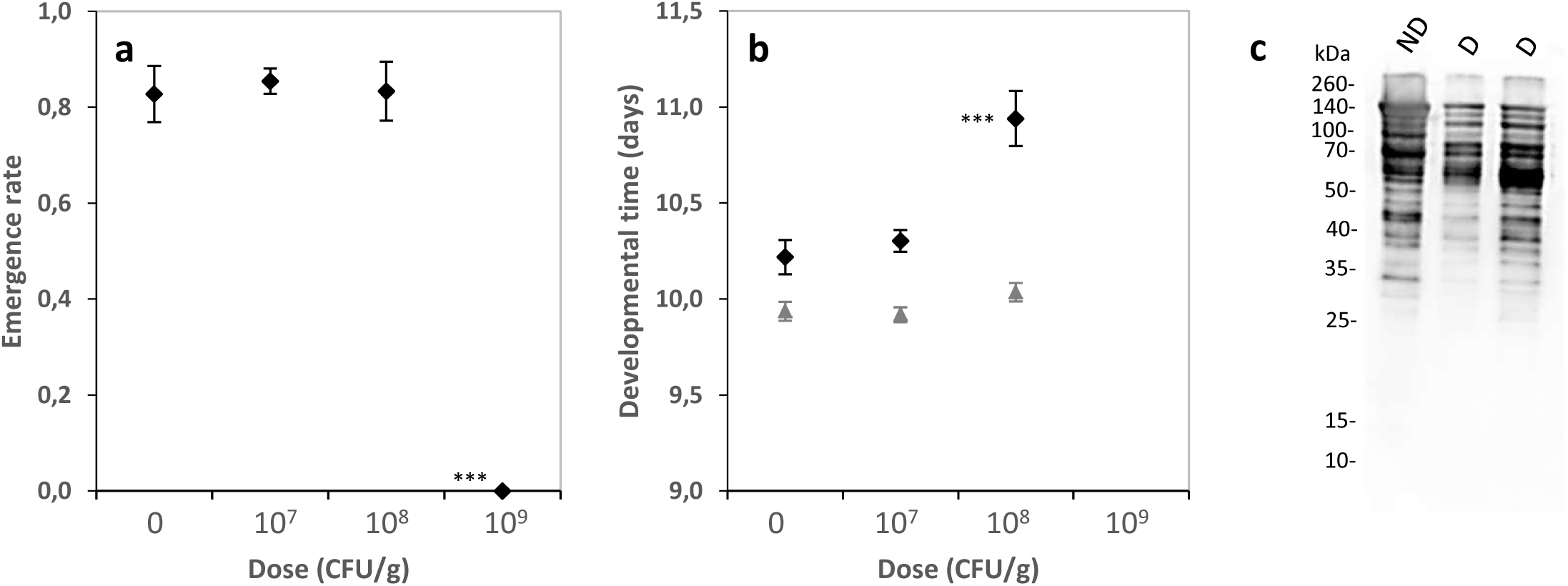
Evaluation of the role of small molecular weight components of *Btk* Delfin^®^ A (dialysis; membrane cut-off: 8-10 kDa) in the altered development of *D. melanogaster* Canton S. (**a**) Emergence rate (mean ± s.e.m.) and (**b**) developmental time (mean ± s.e.m.) on increasing doses of dialyzed Delfin^®^ A. *N* = 3 experiments of 4 replicates with 20 eggs per dose for the emergence rate, *N* = 2 experiments of 4 replicates per dose for the developmental time. Results of *post hoc* comparisons of each dose with the control: · 0.05<*P*<0.1; * 0.01<*P*<0.05; ** 0.001<*P*<0.01; *** *P*<0.001. (**c**) Anti-Cry1A probed immunoblot of non-dialyzed (ND) and dialyzed (D) suspensions showing the decrease in the amount of ∼ 130 kDa protoxins and the increase in that of ∼ 60/70 kDa activated toxins after dialysis.

Cry1A profiles of dialyzed Delfin^®^ A suspensions, like those of the non-dialyzed ones, comprised 130-kDa pro-toxins and 60-70 kDa activated toxins, but also showed toxin degradation as evidenced by additional smaller fragments of activated toxins (Fig. 7c). The respective roles of *Btk* toxin fragments and spores in the alterations of *D. melanogaster* development were further explored through experiments of dialysis such as those previously described, followed by successive centrifugations to eliminate most of the spores and toxin crystals. Despite variation between experiments, ER was strongly affected only in one of the three experiments while DT was always significantly increased when flies were reared in presence of centrifuged supernatants that contained a limited range of Cry 1A toxin fragments (Supplementary information S6).

## Discussion

The increasing use of bioinsecticides based on *Bacillus thuringiensis* (*Bt*) raises concern about their potential non-intentional side-effects on non-target organisms, and biodiversity in general, due to their partially specific targeting,^[33,34,71]^ persistence in the environment,^[35,36,38,40,41,44,72]^ and the requirement of and recommendations for repeated spraying to reach the desired pest control level.^[43]^ Especially, side-effects of chronic exposure on non-target organisms, including insects present on treated areas, remain under-evaluated. Here, we have tested the side-effects of ingestion of *Bt* bioinsecticide formulations (mainly made of *Bt kurstakii* strains (*Btk*) but also of *Bt israelensis* (*Bti*)) throughout the entire development of several non-target species of *Drosophila* flies which are naturally present in treated areas. While formulation doses up to those recommended for each field sprayings (≤ 10^6^ CFU/g of medium) had no effect on *Drosophila* development, mortality and/or developmental delay occurred markedly from doses only 10 times and 50 times higher than the maximum recommended dose of the main tested *Btk* formulation for *D. suzukii* (10^7^ CFU/g) and the *D. melanogaster* strains (5×10^7^ CFU/g), respectively. Development alterations were already strong at these doses, suggesting the occurrence of alterations already at lower doses, i.e. between 10^6^ and 10^7^ CFU/g for *D. suzukii*, and between 10^7^ and 5×10^7^ CFU/g for *D. melanogaster* strains. Further analyses would be needed to verify this possibility. Besides, all the tested species, except *D. simulans*, were strongly affected at 10^8^ CFU/g, and no (or extremely limited) fly development occurred at the highest tested dose (10^9^ CFU/g), equivalent to 1000 times the maximum recommended dose but far below common acute intoxication doses used in many studies.^[5]^ Recommended doses are those for each spraying on a homogeneous and dry zone without covering areas. In the field, recommended repeated sprayings of stabilized formulation and rainfall washouts following spraying may increase *Bt* spores and toxins presence in both space and time. While the highest dose tested here (10^9^ CFU/g) would be hardly reached in the field, the minimal doses at which the fly development was markedly impacted and the lower ones from which changes in development appeared may possibly be obtained. Furthermore, the minimal quantity of *Bt* formulation inducing development alterations may be even lower since a single *Drosophila* larva is unlikely to process 1g of medium given its size and feeding rate. Our data also evidenced a window of *Btk* susceptibility during larval development, with the ingestion during the 1^st^ larval instar accounting for a large part of the observed detrimental effects on the development (see below in the discussion).

When testing for non-intentional generic effects of *Bt* formulations, similar patterns of development alterations were observed but shifted to higher doses with two other *Btk* formulations and a formulation of *Bti*: there was no effect on *D. melanogaster* development at the doses up to 10^8^ CFU/g but a strong detrimental effect at the highest dose tested, 10^9^ CFU/g. The three *Btk* formulations tested, based on two different bacterial strains (see Methods), had similar profiles of Cry1A protoxins and activated toxins, but they differed in their efficient spore contents, formulation type, and likely additives, which may account for the observed variation in the half-lethal dose. The *Bti* formulation, widely used against Dipteran Nematocera insects (e.g. mosquitoes, black flies),^[73]^ impacted *D. melanogaster* development only at the highest dose tested. The impacts of *Btk* formulations on *D. melanogaster* development are consistent with growing evidence suggesting a partly specific targeting of *Bt.*^[33,74]^ Until recently, it has been generally accepted that the mode of action of *Bt* after ingestion by insects relies on key steps of specific binding of proteolyzed *Bt* toxins to receptors of midgut epithelial cells, defining targets for each *Bt* subspecies.^[15,19,22]^ Several primary and secondary types of toxin receptors, including cadherin-like proteins, aminopeptidases, GPI-anchored alkaline phosphatases,^[10]^ and more recently the ATP dependent binding cassette reporter C2,^|75]^ have been identified in Lepidoptera and Diptera mosquitoes. Focusing on the action of *Btk* targeting Lepidoptera, no Lepidoptera cadherin-like Cry receptor orthologues were found in *Drosophila*,^[75]^ supporting the idea that these flies would not be affected by the spraying of *Btk* formulation. The existence of other types of Cry receptors in *Drosophila* flies would explain the observed developmental impacts but remains to be investigated. In addition, the substantial amounts of active Cry1A toxin fragments in *Btk* formulations could compensate for the possible lack of solubilization of protoxin crystals in the fly midgut and proteolytic activation of toxins by fly gut proteases, both required for Cry activity in insect larvae.^[19]^ Other toxins synthesized by *Btk* and present in the formulations could also play a role in the observed cross-order activity as some, such as Cry2A, have an insecticidal effect on both Lepidoptera and Diptera.^[76]^

The lack of effect of ingestion of *Bacillus subtilis* or *Btk* Cry-free 4D22 on the development of *D. melanogaster* excludes that development alterations result from severe disruption of digestion and nutrient uptake/competition in the presence of high spore/bacteria loads in the larval gut throughout development. It supports the idea of a synergistic action of *Btk* spores and Cry toxins, consistent with the models of *Bt* action on insect larvae in which toxins first breach the gut epithelium, allowing the gut content, including *Bt* spores, to colonize the hemocoel.^[19,22-24]^ The partially reproduced mortality rate and delayed development in dialysis experiments may further indicate that low diffusible molecular weight compounds in *Btk* formulations (e.g., culture media residues, salts, additives) may contribute to these development alterations. This is supported by the lack of impact on *D. melanogaster* development of the ingestion of laboratory-produced spores and Cry toxins of a *Btk* 4D1 strain (or HD1, a reference strain used as a control strain here, not used in commercial formulations) used without additives, even at the highest dose 10^9^ CFU/g (additional information S7; Fig. S7a, b). The *Btk* 4D1 culture contained few active Cry toxins and smaller toxin fragments, in contrast to commercial *Btk* formulations (Fig. S7e), supporting the possible contribution of these toxin fragments to the cross-order activity of *Btk* formulations on *Drosophila*.

As previously reported for *D. suzukii* exposed to laboratory-produced *Btk* cultures,^[53]^ mortality of *D. melanogaster* during development on *Btk* formulation occurred early in development. First and second instars larvae are likely highly exposed due to their high feeding rate and their exponential growth.^[77]^ As the observed larval mortality was only about 40% at the highest dose (10^9^ CFU/g) (Figure 2), while none of the individuals reached the pupal stage at this dose (Figure 4), the remaining mortality likely occurred during the third larval stage, maybe due to a delayed action of *Btk* spores and toxins. Interestingly, alterations of the development (mortality and delayed emergence) mimicked those typically generated by nutritional stress conditions in insect larvae.^[78,79]^ Accordingly, the development alterations were partially rescued on a protein rich fly medium, probably through compensatory protein intake, as in other arthropod species.^[79-81]^ In addition, the sex ratio of flies was strongly biased towards males after development on the dose of *Btk* formulation affecting fly emergence (10^8^ CFU/g) and under low protein conditions. This highlights the importance of nutritional conditions in *Btk* impacts on *Drosophila* development, with sex-specific differences in larval susceptibility to environmental stressors, here the accumulation of *Btk* formulation, under protein restriction conditions as reported previously in *D. melanogaster.*^[82]^

The development on sublethal doses of *Btk* formulation did not dramatically affect the longevity of *D. melanogaster* adults and the lifetime offspring number. Developmental exposure to *Btk* doses that slightly and strongly reduced the likelihood of reaching the adult stage even gave to males and females a dose-dependent longevity benefit, in addition to the male higher longevity observed in mixed-sex populations,^[83]^ and slightly increased the offspring number (although not significantly). Surviving the exposure to *Btk* formulation throughout the development has likely selected for fitter individuals. This is similar to the increased longevity of adult insects that have survived developmental nutritional stress,^[84,85]^ or are resistant to environmental stressors.^[83]^

The origin of *Drosophila* (species and population/strain) influenced the magnitude of the development alterations induced by *Btk* formulation. For *D. melanogaster*, all the strains tested were susceptible to *Btk* formulation with both mortality and delayed development at the same dose, but with variation in the effect magnitudes. This suggests potential population-specific differences in susceptibility to *Btk* formulation accumulation in the environment, and hence potential spatial and temporal heterogeneity of *Btk* spraying impacts for each *Drosophila* species. Among the other seven species tested, differences in susceptibility to *Btk* formulation, in terms of effect magnitude and type of development alteration (mortality and/or developmental delay), occurred between *Drosophila* species, regardless of their phylogenetic distances. For the *Drosophila* subgenus, *D. simulans* was less susceptible than its sister species *D. melanogaster*, whereas the African *D. yakuba* experienced similar impacts on the development as *D. melanogaster*. The three species *D. immigrans, D. subobscura* and *D. hydei* were similarly more susceptible than *D. melanogaster*, but with slight differences in effect magnitudes. The phylogenetically distant *D. busckii* (*Dorsilopha* subgenus) was the least affected of all the species tested in terms of developmental mortality, but its development was strongly delayed. The five species *D. melanogaster, D. simulans, D. hydei, D. immigrans*, and *D. busckii* belong to the guild of cosmopolitan domestic *Drosophila* species, *D. subobscura* is sub-cosmopolitan species, and *D. busckii* is an opportunistic frugivorous species.^[86]^ All these species coexist frequently and compete on the same discrete and ephemeral rotting fruit patches, with seasonal variations in the composition of the fly community.^[56-59]^ At the community level, differences in the species susceptibility to accumulation of *Btk* formulation could modify larval competition conditions and lead to additional local and temporal variations in *Drosophila* communities’ composition. The potential side-effects of *Bt* sprays on non-target *Drosophila* communities would be hardly predictable as they would depend on spatial patterns of *Bt* accumulation. A formal mesocosm study of *Drosophila* community dynamics under exposure to *Btk* formulation, at least under semi-field conditions, would allow assessing the consequences of *Bt* accumulation on species competition and community composition.

As for the other species, the presence of *Btk* formulation impacts the development of the invasive *D. suzukii*, as recently reported by Cossentine et al.,^[53]^ this species being the most susceptible here with effects already clearly detectable at 10 times the recommended spraying dose. Compared with the other seven species that live on rotten fruits, *D. suzukii* poses a threat to fruit production because it feeds and lays eggs in healthy ripening soft fruits^[87-89]^ and hence colonizes orchards and vineyards earlier during the fruit season. Despite its oviposition mode with a saw-like ovipositor inserted inside ripening soft fruits,^[87]^ the exposure of *D. suzukii* larvae may be quite similar to that of larvae of other *Drosophila* species laying on the surface of fermenting fruits or rotting fruit flesh. Indeed, the saw-like ovipositor likely carries *Bt* spores and toxins from the surface of treated fruits when piercing the fruit skin, and additional *Bt* may then enter and germinate in the fruit through the drilled holes. From an ecological point of view, the greater susceptibility of *D. suzukii* to the accumulation of *Btk* formulation in the environment might mitigate the potential ecological burden of its invasion for local communities of *Drosophila* frugivorous species in orchards. Alternatively, as *D. suzukii* attacks on fruits can accelerate their decomposition by microorganisms, its higher susceptibility to *Btk* could reduce the number of fruits made suitable for other *Drosophila* species.

In conclusion, our results showed that at recommended *Bt* doses, no visible effects on *Drosophila* was observed, but accumulation of *Btk* biopesticide above these doses can potentially impact these non-target insects. The magnitude of this impact possibly depends on both the formulation used and the fly species. Although our study was carried out under controlled laboratory conditions which may dramatically differ from natural conditions encountered in the field (temperature, pH, humidity, food availability, photoperiod, predator/parasite/pathogen pressures, etc…), standard laboratory strains and flies derived from wild populations recently collected exhibited similar patterns of development alterations, suggesting our results may not be specific to laboratory-influenced genetic backgrounds. Recent studies have reported similar adverse side-effects due to repeated sprayings of the *Bti* formulation, directly on non-target organisms,^[31]^ and indirectly on predators via food webs.^[90]^ These studies and the data presented here highlight that pest control with *Bt* bioinsecticides should be done with caution in the field to avoid, or at least limit, potential non-intentional side-effects on non-target organisms and hence on biodiversity. At last, *D. melanogaster*, a model species in many research fields, could also serve as a study model to identify the mechanisms underlying these side-effects and/or the potential emergence of resistance to these biopesticides.

## Acknowledgements

We thank Xiao Han, Jingru Li and Abir Oueslati for help with preliminary experiments, L. Kremmer, C. Rebuf and O. Magliano for providing and rearing flies and help in preparing fly medium, A. Brun-Barale for the production of *Bacillus subtilis* spores, D. Pauron for preparation of Cry1A toxin, Hugo Mathé-Hubert for advice on statistical analyses, and M. Amichot for helpful discussions. The Cry1A antibody was produced in collaboration with the INRA-PFIE platform (Nouzilly, France).

## Financial Supports

This work was supported by the French National Agency for Research (ANR-13-CESA-0003-001 ImBio), the European Union’s Seventh Framework Program for research, technological development and demonstration under grant agreement No. 613678 (DROPSA), the “Investments for the Future” LABEX SIGNALIFE (ANR-11-LABX-0028), the INRA Plant Health Department (to MPNE and JLG), the CNRS (to AG), and the University Nice Côte d’Azur (to MP).

## Author contributions

AB, MPNE, AG, JLG and MP designed the experiments. AB performed the experiments with contributions of MPNE. AB performed the statistical analyses. AB, JLG, and MP wrote the manuscript with contributions from all the authors.

## Additional information

**Supplementary information**

