## Supplementary Information for "Differential side-effects of *Bacillus thuringiensis* bioinsecticide on non-target *Drosophila* flies"

### Contents

S1. Equivalence of tested doses of commercial formulations and spore production in CFU/cm$^2$.

S2. Sex-ratio of *Drosophila melanogaster* Canton S flies on *Bt* commercial formulations.

S3. Sex-ratio of four *D. melanogaster* strains on *Btk* formulation.

S4. pH of the fly medium in the presence of increasing doses of *Bt* formulations.

S5. Sex-ratio of seven *Drosophila* species on *Btk* formulation.

S6. Role of toxin fragments and spores of *Btk* formulations in the alterations of the development of *D. melanogaster* Canton S.

S7. Development of *D. melanogaster* Canton S on *Btk* 4D1 (or HD1).

### S1. Conversion of tested doses of commercial formulations from CFU/g of fly medium used in the experiments to CFU/cm$^2$.

Applications of *Bt* formulations in the field follow the dose recommendations of the manufacturers, expressed in CFU per surface unit. Since in our experiments, fly larvae forage not only on the surface of the fly medium but also within the medium, we mixed formulations and spore productions with the medium and used doses expressed as CFU per quantity of food, i.e. CFU/g of medium. To facilitate comparison of the doses tested with those recommended by the producers, we provide here the recommended average doses for the field application of each formulation used (Table S1a), and the approximate conversion doses calculated in CFU/cm$^2$ of our doses for each type of vial used in our experiments (Table S1b).

**Table S1a. Recommended field application doses.** Dose range (min, max) and average dose recommended by manufacturers in CFU/cm$^2$ for each of the four *Bt* formulations we used.

| ***Bt* formulation** | **Min** | **Max** | **Mean** |
|---|---|---|---|
| Delfin® A | $7.5\times10^4$ | $7.5\times10^5$ | $4.1\times10^5$ |
| Delfin® B | $3.75\times10^4$ | $3.75\times10^5$ | $2.1\times10^5$ |
| Scutello DF | $2.2\times10^4$ | $2.2\times10^5$ | $1.2\times10^5$ |
| Vectobac® WG | $7.5\times10^4$ | $6\times10^5$ | $3.4\times10^5$ |

**Table S1b. Approximate calculated conversion of tested doses in CFU/cm$^2$.** Conversion of the doses used for the dose-response assays, expressed in CFU/g, into doses expressed in CFU/cm$^2$, based on the surface and the quantity of fly medium in each type of vial used in the experiments.

| **Features of vials used in the different tests** | | | | | |
|---|---|---|---|---|---|
| Diameter (cm) | | 3 | 3.3 | 4.6 | |
| Surface (cm$^2$) | | 7 | 8.5 | 16 | |
| Provided fly medium quantity (g) | | 1 | 2 | 6 | |
| Fly medium distribution (g/cm$^2$) | | 0.14 | 0.24 | 0.37 | |
| **Used doses in CFU/g** | | **Dose equivalence in CFU/cm$^2$ per tube** | | | **Mean** |
| Lowest dose | $1\times10^5$ | $1.4\times10^4$ | $2.4\times10^4$ | $3.7\times10^4$ | $2.5\times10^4$ |
| | $1\times10^6$ | $1.4\times10^5$ | $2.4\times10^5$ | $3.7\times10^5$ | $2.5\times10^5$ |
| | $1\times10^7$ | $1.4\times10^6$ | $2.4\times10^6$ | $3.7\times10^6$ | $2.5\times10^6$ |
| | $1\times10^8$ | $1.4\times10^7$ | $2.4\times10^7$ | $3.7\times10^7$ | $2.5\times10^7$ |
| Highest dose | $1\times10^9$ | $1.4\times10^8$ | $2.4\times10^8$ | $3.7\times10^8$ | $2.5\times10^8$ |

**S2. Sex-ratio of *Drosophila melanogaster* Canton S flies on *Btk* commercial formulations.**

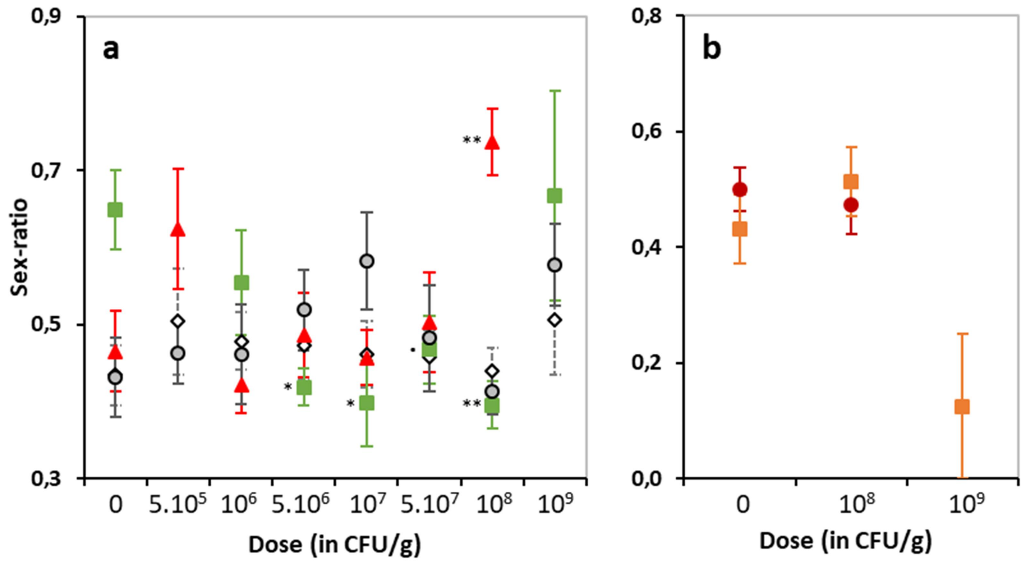


**Figure S2. Fly sex-ratio of *D. melanogaster* Canton S after development on increasing doses of *Bt* and *B. subtilis*.** (**a**) Sex-ratio (expressed as the male proportion; mean ± sem) after complete development of 20 initial eggs on a low protein/high-sugar medium containing increasing doses of *Btk* formulation Delfin® A (red triangles), control Cry-free *Btk* 4D22 strain (open lozenges), the mosquito-targeting *Bti* Vectobac® (green squares), and the non-pathogenic *Bacillus subtilis* (grey circles). For Vectobac® and *B. subtilis*, *N* = 4-7 per dose; for Delfin® A and *Btk* 4D22, *N* = 9-12 for the control, *N* = 3 for $5.10^5$ and $10^9$, *N* = 4-9 for $10^6$, *N* = 7-14 from $5.10^6$ to $10^8$. (**b**) Sex-ratio on increasing doses of the two other *Btk* formulations: Delfin® B (dark red circles) and Scutello DF (orange squares). *N* = 4 replicates of 20 eggs per dose and formulation, except for controls and $10^8$ CFU/g of Delfin® B (9-10 replicates of 20 eggs). Missing data points correspond to null emergence rates. Asterisks show results of *post hoc* tests comparing each dose to the control: · $0.05<p<0.1$; * $0.01<p<0.05$; ** $0.001<p<0.01$ (see Table S1).

**Table S2.** Results of statistical analyses to assess the effect of the dose of formulation/spore production and interaction with treatment on the sex-ratio of *D. melanogaster* Canton S flies. See significant *post hoc* comparisons of the doses with the control Fig. S2.

| Source of variation/Data | $\chi^2$ | d.f. | *P* value |
|---|---|---|---|
| **Development on *Btk* Delfin® A, *Btk* 4D22, *Bti* Vectobac®, *Bacillus subtilis*** | | | |
| Dose × Treatment | 31.8 | 19 | 0.033 |
| Dose for each treatment: | | | |
| - Delfin® A | 15.6 | 6 | **0.016** |
| - 4D22 | 2.19 | 7 | 0.95 |
| - Vectobac® | 11.9 | 6 | 0.064 |
| - *B. subtilis* | 6.05 | 7 | 0.53 |
| **Development on *Btk* Delfin® B and Scutello DF (dose effect)** | | | |
| - Delfin® B | 0.077 | 1 | 0.78 |
| - Scutello DF | 2.48 | 2 | 0.29 |

S3. Sex-ratio of four *D. melanogaster* strains on *Btk* formulation.

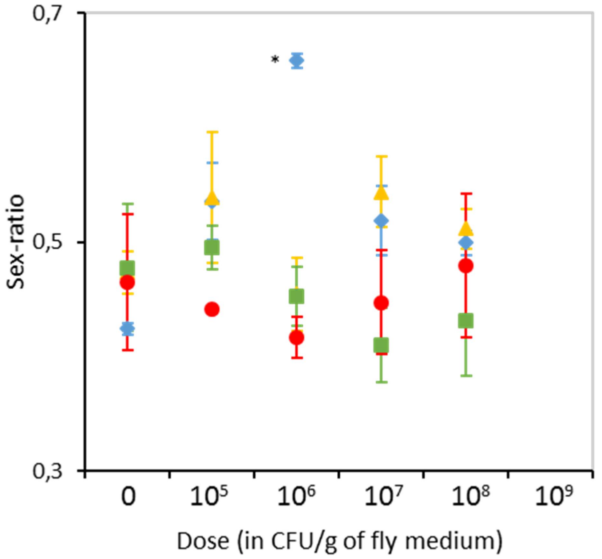


**Figure S3. Fly sex-ratio of four *D. melanogaster* strains after development on increasing doses of the *Btk* Delfin® A.** Sex ratio expressed as the male proportion (mean ± sem) of *D. melanogaster* flies of the wild-type strains Canton S (blue lozenges), Nasrallah (strain 1333; yellow triangles), and Sefra (green squares), and the double-mutant strain YW1118 (red circles) after complete development on a high-protein/sugar-free medium. $N = 4$ replicates of 50 initial eggs per dose and fly strain. Asterisks show the results of *post hoc* tests comparing each dose to the control: * $0.01<p<0.05$.

**Table S3.** Results of statistical analyses to assess the effect of the *Btk* Delfin® A dose on the sex-ratio of flies for the four *D. melanogaster* strains. See significant *post hoc* comparisons of the doses with the control in Fig. S5.

| Data | $\chi^2$ | d.f. | *P* value |
|---|---|---|---|
| - Canton S | 9.15 | 4 | 0.057 |
| - Nasrallah | 2.06 | 4 | 0.73 |
| - Sefra | 1.52 | 4 | 0.82 |
| - YW1118 | 0.81 | 4 | 0.94 |

**S4. pH of the fly medium in presence of increasing doses of *Bt* formulations.**

*Bt* formulations are buffered at low pH (*Bt* activity being stable in a 3-11 range of pH; Brar *et al.*, 2006). In addition, changes in the medium pH may affect *Drosophila* development (Hodge and Caslaw 1998; Hodge 2001). To control for a confounding effect of low formulation pH in the development alterations, we measured the pH of the medium mixed with increasing doses of *Btk* formulations after fly emergence.

Methods

pH was measured by soaking a piece of pH paper onto the surface of the fly medium, for the same dose range of Delfin® A, *Btk* 4D22, Delfin® B, and Scutello DF as tested in the dose-response on the development-related traits.

Results

The pH varied between 4.9 and 6.7 regardless of the presence and the dose of the three *Btk* formulations and of the Cry-free *Btk* 4D22 strain.

### S5. Sex-ratio of seven other *Drosophila* species on *Btk* formulation.

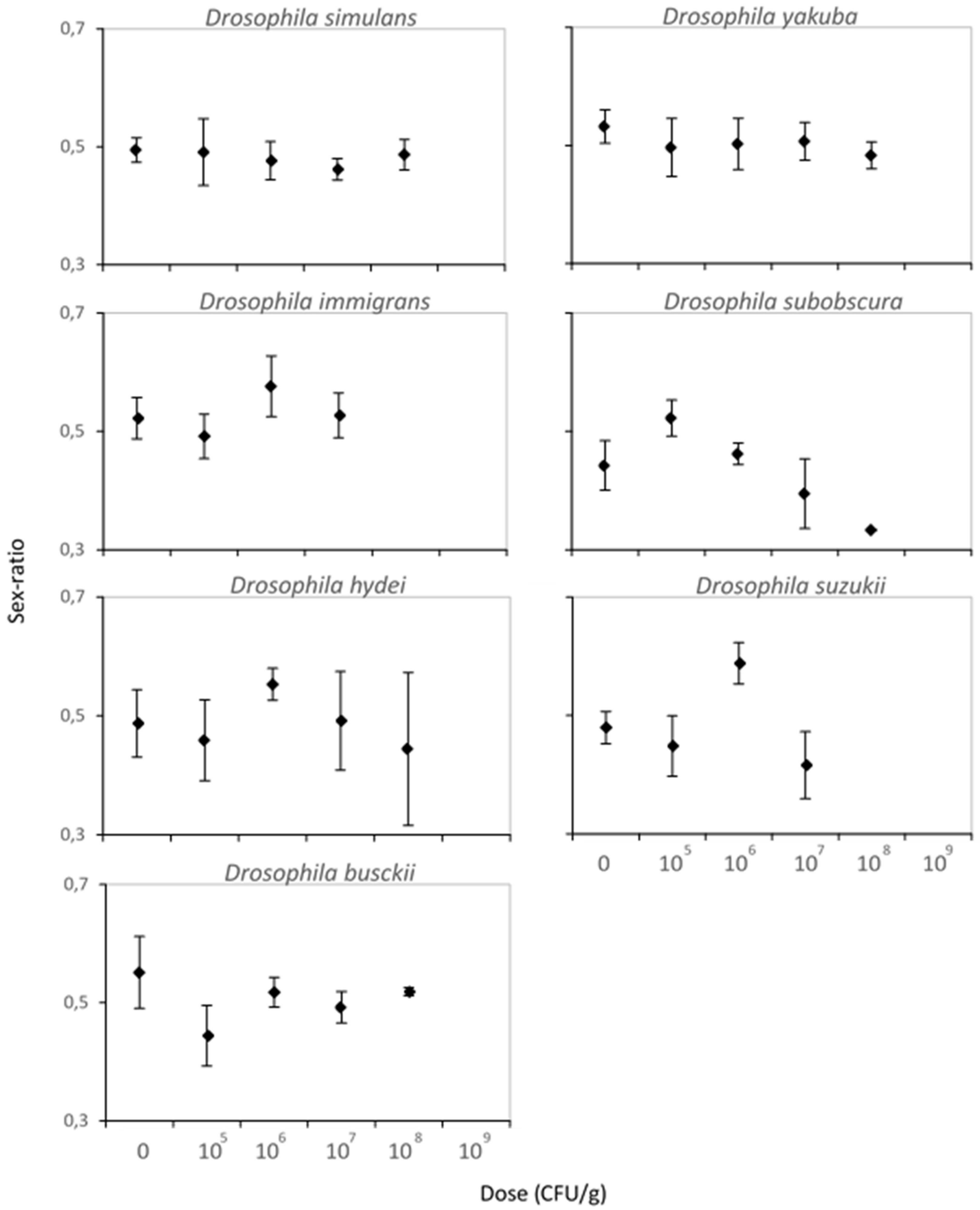


**Figure S5. Fly sex-ratio of seven *Drosophila* species after development on increasing doses of *Btk* Delfin® A.** Sex ratio expressed as the male proportion (mean ± sem) after complete development of the *D*rosophila species on a high-protein/sugar-free medium. $N = 4$ replicates of 50 initial eggs per dose and species (or 30 initial eggs for *D. hydei*, *D. suzukii* and *D. immigrans*). No statistical difference was observed with *post hoc* tests comparing each dose to the control (see also Table S5).

**Table S5.** Results of statistical analyses to assess the effect of the Delfin® A dose on the sex-ratio of flies for each of the 7 additional *Drosophila* species.

| Data | $\chi^2$ | d.f. | *P* value |
|---|---|---|---|
| - *D. simulans* | 0.44 | 4 | 0.98 |
| - *D. yakuba* | 0.26 | 4 | 0.87 |
| - *D. hydei* | 1.98 | 4 | 0.74 |
| - *D. immigrans* | 1.44 | 3 | 0.70 |
| - *D. subobscura* | 6.95 | 4 | 0.14 |
| - *D. suzukii* | 5.33 | 3 | 0.15 |
| - *D. busckii* | 3.92 | 4 | 0.42 |

**S6. Role of toxin fragments and spores of *Btk* formulation in the alteration of the development of *Drosophila melanogaster* Canton S.**

In addition to the dialysis experiment, the respective roles of *Btk* toxin fragments and spores was explored preliminarily with successive centrifugations of a dialyzed suspension of Delfin® A at increasing speeds (1,000 g, 5,000 g, and 15,000 g).

Methods

Dialysis was performed on a suspension of $2\times10^{10}$ CFU of Delfin® A against PBS ($KH_2PO_4$ 1.06 mM, $Na_2HPO_4(2H_2O)$ 3mM, NaCl 154 mM, qsp distilled water) at 250 rpm and 4°C overnight, using an 8-10 kDa MW cut-off membrane (ZelluTrans, Roth®). The dialyzed suspension was centrifuged at 1,000 g (10 min, 4°C). The supernatant was collected carefully and centrifuged successively at 5,000 g and 15,000 g (10 min, 4°C). The CFUs of the supernatants collected after each centrifugation were estimated as described in the main methods. Effects of the three supernatants were tested on egg-to-adult development, by measuring the emergence rate (ER) and the development time (DT) on the low-protein/high-sugar fly medium. The experiment was repeated three times. Each of the supernatants was also analyzed on a 12.5 % SDS-PAGE and compared to the dialyzed suspension for the presence of Cry1A pro-toxins, activated toxins and toxin fragments by Western-blot using an in-house anti-Cry1A rabbit polyclonal antibody.

Results

After centrifugation, the three supernatants contained few spores, corresponding to doses below those inducing development alterations (mean CFUs ± sem per 100 µl: $6.4\times10^5 \pm 0.5$ after 1,000 g, $1.8\times10^4 \pm 0.01$ after 5,000 g, $8.9\times10^3 \pm 1.1$ after 15,000 g). The supernatant mixed with the low-protein/high-sugar fly medium provided an equivalent to the formulation dose $5\times10^8$ CFU/g at which development alterations were previously observed (see Figure 1a in the main results section).

ER on the supernatants decreased by up to 60% compared to the buffer control, but only in one of the three experiments (Fig. S2a, Table S2). By contrast, despite large between-experiments variation, DT increased on the three supernatants by minimum ~10% (~1 day) compared to the buffer control in each of the three experiments (Fig. S2b, Table S2). The Cry1A profile showed that the centrifugation at 1,000g eliminated almost all of the ~130-kDa pro-toxins and all of the ~60-70 kDa activated toxins, while smaller fragments of ~40 kDa remained in the three supernatants (Fig. S2c).

Interestingly, without prior dialysis of the suspension of Delfin® A, the three supernatants of successive centrifugations prevented fly development at a dose corresponding to $5\times10^8$ CFU/g (no individual reached the pupation; data not shown).

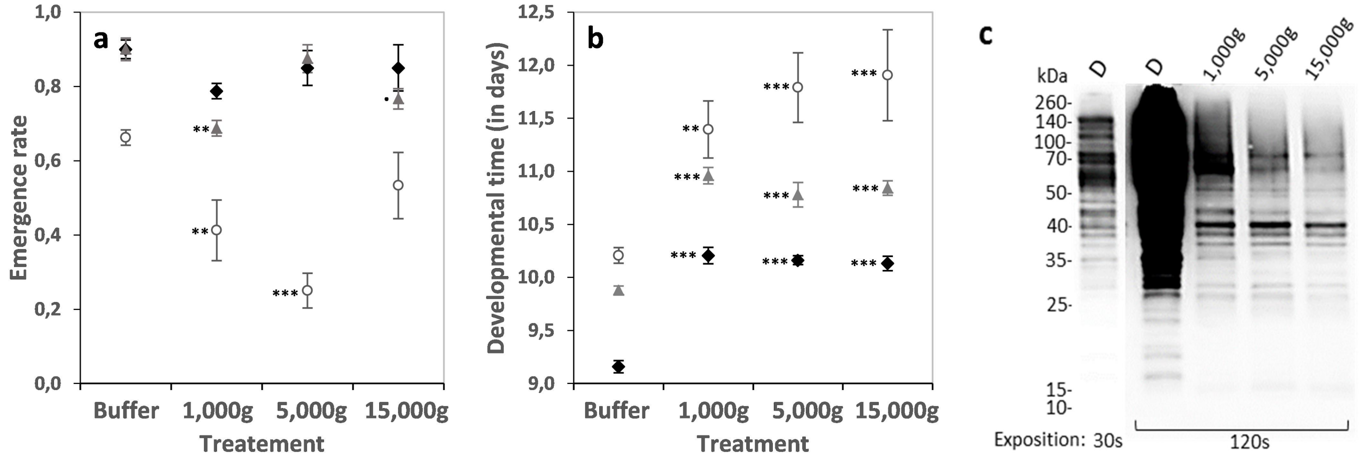


**Figure S6. Effect of the spores and toxin fragments of *Btk* Delfin® A in the alterations of *D. melanogaster* Canton S development with dialysis and centrifugation.**

(**a**, **b**) Emergence rate and developmental time (mean ± sem) in presence of the successive centrifugation supernatants (1,000 g, 5,000 g, and 15,000 g) of a dialyzed (D) suspension of Delfin® A. $N = 3$ experiments of 3-4 replicates of 20 eggs per treatment. Open circles symbols correspond to experiment 1, black lozenges to experiment 2, and grey triangles to experiment 3. Asterisks show the results of *post hoc* tests comparing each centrifugation treatment to the control (Buffer): • $0.05<p<0.1$; ** $0.001<p<0.01$; *** $p<0.001$ (see Table S2). (**c**) Profile of anti-Cry1A immunoblotting of a dialyzed suspension of Delfin® A (D) at two exposure times (30s, 120s), and of supernatants from the three centrifugation speeds from experiment 2.

**Table S6.** Results of statistical analyses to assess the effect of the centrifugation treatment on the emergence rate and the developmental time in three experimental blocks. See significant *post hoc* comparisons of the doses with the control in Fig. S3.

| Data | $\chi^2$ | d.f. | *P* value |
|---|---|---|---|
| <u>*Emergence rate*</u> | 30.3 | 3 | **< 0.0001** |
| - experiment 1 | 31.6 | 3 | **< 0.0001** |
| - experiment 2 | 3.94 | 3 | 0.27 |
| - experiment 3 | 14.9 | 3 | **0.0019** |
| <u>*Developmental time*</u> | 44.7 | 3 | **< 0.0001** |
| - experiment 1 | 15.9 | 3 | **0.0012** |
| - experiment 2 | 41.7 | 3 | **< 0.0001** |
| - experiment 3 | 34.2 | 3 | **< 0.0001** |

**<u>S7. Development of *D. melanogaster* Canton S on *Btk* 4D1 (or HD1).</u>**

The standard wild-type strain Canton S was reared on low-protein/high-sugar fly medium mixed with laboratory-produced spores and toxins of *Bt kurstaki* reference strain 4D1 (or HD1, a reference strain used as a control strain here, not used in the commercial formulations). This strain produces *Btk* Cry toxins Cry1Aa, Cry1Ab, Cry1Ac, Cry2Aa, Cry2Ab, and spores. The laboratory production does not contain additives or vegetative cells (eliminated), and Cry 1A toxins are still mainly in the form of protoxins and few as activated 60-70 kDa toxins (no multiple smaller Cry fragments).

<u>Methods</u>

*Btk* 4D1 culture was produced as described for the acrystillipherous strain 4D22 in the main Methods. Number of CFUs was estimated to reach the desired dose. As for *Btk* formulation Delfin® A and *Btk* 4D22, tested doses ranged from $5\times10^5$ CFU/g to $10^9$ CFU/g by mixing 100 µl of *Btk* 4D1 suspension for 1 g of low-protein/high sugar fly medium.

As described in the main Methods, eggs of Canton S flies were collected after mass oviposition. Replicate vials of 20 eggs for 2 g of fly medium were incubated under standard laboratory conditions (25°C, 60% relative humidity, light-dark cycle 12:12). Emergence rate, developmental time and sex-ratio were calculated as described in the main Methods.

<u>Results</u>

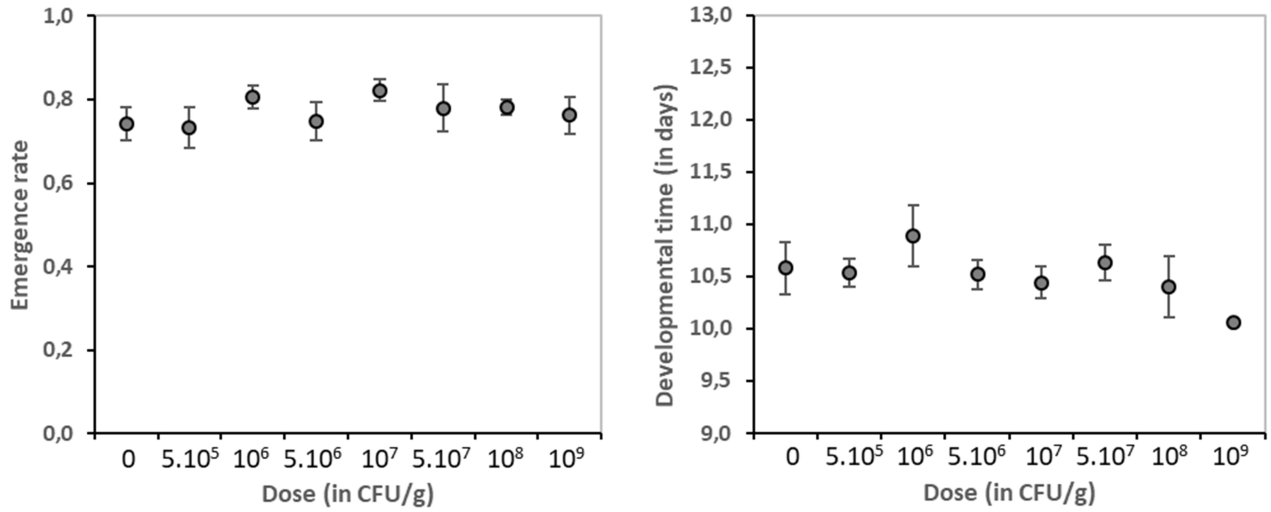


**Figure S7a. Development of *D. melanogaster* Canton S on increasing doses of *Btk* 4D1.** Emergence rate (mean ± sem) and developmental time (mean ± sem) on sugar-rich/low-protein fly medium. $N$ = 3-4 of 20 eggs for $5\times10^5$ and $10^9$ CFU/g, $N$ = 9-13 of 20 eggs for control and all the other doses.

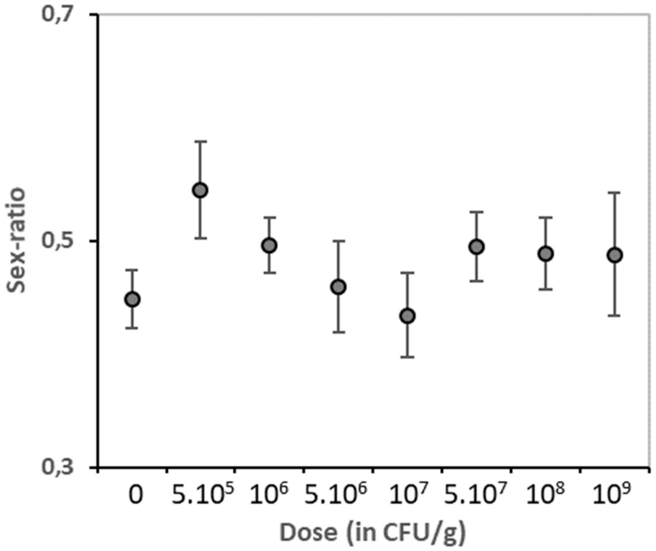


**Figure S7b. Fly sex-ratio (mean ± sem) of *D. melanogaster* Canton S after development on increasing doses of *Btk* 4D1.** *N* = 3-4 of 20 initial eggs for $5 \times 10^5$ and $10^9$ CFU/g, *N* = 9-13 of 20 initial eggs for control and all the other doses.

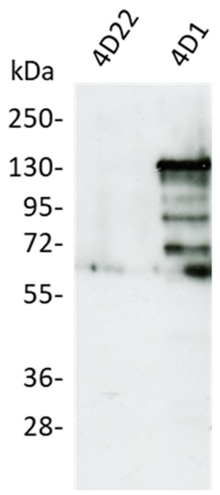


**Figure S7c.** Anti-Cry1A probed immunoblot of laboratory-produced *Btk* 4D22 that does not produce Cry toxins and of laboratory-produced *Btk* 4D1 showing the main presence of ~ 130 kDa protoxins, and the marginal presence of ~ 60/70 kDa activated toxins.
